# Longitudinal dynamics of human B-cell response at single-cell level in response to Tdap vaccination

**DOI:** 10.1101/2021.10.08.463649

**Authors:** Indu Khatri, Annieck M. Diks, Erik B. van den Akker, Liesbeth E.M. Oosten, Jaap Jan Zwaginga, Marcel J.T. Reinders, Jacques J.M. van Dongen, Magdalena A. Berkowska

## Abstract

Adaptation of the immune system to mount an adequate immune response against pathogens is a crucial function of the adaptive immune system. To better characterize a successful vaccination response, we performed longitudinal (days 0, 5, 7, 10, and 14 after Boostrix vaccination) analysis of the single cell transcriptome as well as the B-cell receptor (BCR) repertoire (scBCR-rep) in plasma cells of an immunized donor and compared it with baseline B cell characteristics as well as flow cytometry findings.

Based on the flow cytometry knowledge and literature findings, we discriminated individual B cell subsets in the transcriptomics data and traced over-time maturation of plasmablasts/plasma cells (PB/PCs) and identified the pathways associated with the plasma cell maturation. We observed that the repertoire in PB/PCs differed from the baseline B cell repertoire e.g. regarding expansion of unique clones in post-vaccination visits, high usage of *IGHG1* in expanded clones, increased class switching events post-vaccination represented by clonotypes spanning multiple *IGHC* classes and positive selection of CDR3 sequences over time. Importantly, the *Variable* gene family-based clustering of BCRs represented a similar measure as the gene-based clustering, however, certainly improved the clustering of BCRs, as BCRs from duplicated *Variable* gene families could be clustered together. Finally, we developed a query tool to dissect the immune response to the components of Boostrix vaccine. Using this tool, we could identify the BCRs related to anti-tetanus and anti-pertussis toxoid.

Collectively, we developed a workflow which allows description of key features of an ongoing immune response, such as activation of PB/PCs, Ig class switching, somatic hypermutation, and clonal expansion, all of which are hallmarks of antigen exposure.

## Introduction

The adaptation of the immune system to recognize antigenically distinct epitopes of pathogens is a critical feature of the antibody (Ab) -mediated immunity in response to infection/vaccination. Abs are products of terminally differentiated B cells, plasma cells (PCs), and are the secretory form of B-cell receptors (BCRs). Generation of a highly diverse BCR repertoire (BCR-rep) starts during precursor B cell development in bone marrow where developing B cells assemble immunoglobulin heavy and light chain genes in the process of *V(D)J* recombination. B cells with a functional BCR leave the bone marrow and are ready to recognize antigens in the peripheral immune system. Following antigen recognition, the BCR undergoes affinity maturation in the germinal centers (GCs) through introduction of somatic hypermutations (SHM) [1]. Introduction and subsequent selection of SHM are critical steps in raising the appropriate immune response. Typically, the selected SHM accumulate in so called complementarity-determining regions (CDRs) because CDRs are in direct contact with the antigen, while SHM are less frequent in the framework regions (FWR), because FWRs are responsible for maintaining receptor structure. Additionally, B cells optimize their effector functions by exchanging the initial Cμ constant region by other regions located downstream *IGH* loci. This process is known as class-switch recombination (CSR). Memory B cells that have recognized pathogen/antigen circulate through the peripheral lymphoid system differentiate into plasmablasts and re-enter bone marrow to become long-lived Ab-producing PCs.

The immune response to a booster vaccination or re-encounter of the same antigen would lead to the use of pre-existing Abs as the first line of defense [2]. The encounter with an antigen will initiate the process where new Abs are generated from the pool of the existing memory B cells [3,4]. Using flow cytometry, we have recently shown that in booster vaccination settings (Boostrix), an increase in circulating PC numbers, associated with their phenotypical maturation, occurs as early as 5 days post-vaccination, with a peak in expansion and maturation at day 7 post-vaccination [5]. However, the dynamics of the changes in the BCR-rep after Boostrix vaccination in humans are not known yet. Therefore, we performed longitudinal measurements and analysis of single-cell transcriptomics and single-cell BCR-rep (scBCR-rep) in PCs derived from an individual vaccinated with a Tdap Booster vaccine (Boostrix®, GlaxoSmithKline), and compared them with the total BCR-rep at baseline.

As the Boostrix vaccine contains multiple antigens i.e. diphtheria toxoid (DT), tetanus toxoid (TT), three *Bordetella pertussis* proteins i.e. filamentous hemagglutinin (FHA), pertactin (Prn) and pertussis toxoid (PT), and aluminum hydroxide as an adjuvant [6], we obtained receptor repertoire information against all the components of the vaccine. Although it is known that the majority of PCs in the peak of response are Ag-specific [7,8], we aimed to dissect the immune response against each component. This was, however, subjected to the known pool of Abs against the antigens of Boostrix vaccine. Knowing limitations of the available anti-toxoid Abs for toxoids in vaccine, we developed a method to search the known Ab CDR3 sequences for these vaccine components in our data.

Here, we have analyzed the longitudinal single-cell transcriptomics and scBCR-rep data in PCs derived from an individual vaccinated with Boostrix vaccine and compared them with the total B cells and or plasma cells at baseline. Using transcriptomics data, we have identified additional membrane markers for flow cytometry that can be used to identify the B cell subtypes and maturation of plasma cells. With the scBCR-rep data, we have clustered functionally similar receptors and assessed different components of the repertoire for associations between visits and to baseline. Importantly, we found BCRs associated with the vaccine components in scBCR-rep data.

## Methods

### Inclusion criteria and informed consent from the subject

The donor used in this study was a 28-year-old female, who was recruited as one of 10 healthy adults in a bigger study where volunteers were vaccinated with Boostrix and their immune responses were monitored over time by means of flow cytometry (registration number: P16-214, EUDRACT: 2016-002011-18). This study was approved by the Medical Ethics Committee Leiden-Den Haag-Delft. Inclusion and exclusion criteria were previously published [5]. In short, the donor had to be generally healthy, had completed vaccination schedule according to the Dutch National Immunization program (www.rivm.nl/en/national-immunisation-programme) and had no known or suspected exposure to pertussis infection nor vaccination in the past 10 years. She also provided an informed consent prior to the inclusion. EDTA-blood samples for single-cell sequencing were collected at baseline and days 5 (d5), d7, d10 and d14 post-vaccination, which based on the parent study, corresponded to the peak of plasma cell expansion (d5-d10) and contraction of plasma cell response [5] (**Figure S1)**.

### High throughput B cell and plasmablast/plasma cell sorting

Prior to the sort, 78.2 million PBMCs from 5 different time-points were individually (per time-point) stained with an Ab cocktail for surface markers for 30min at RT in the dark (CD19 BV786, CD24 BV650, CD27 BV421, CD38 PerCP Cy5.5), followed by a life/dead staining (Zombie NIR Fixable Viability kit, Biolegend) according to the manufacturer’s protocol. Next, samples were prepared for multiplexing according to the protocols provided on the 10x Genomics website (www.10xgenomics.com). In short, Fc block was added and after a 10 min incubation at 4°C, unique cell hashing Abs (10x Genomics) were added to each sample separately (per time-point). After incubation for 20 min at 4°C, cells were washed three times and resuspended in PBS. Next, all post-vaccination samples were pooled and sorted on a high-speed cell sorter (BD FACSAria III, BD Biosciences at the flow cytometry core facility of LUMC) using a 85μm nozzle and a pressure of 45PSI, while the baseline sample was sorted separately, but in the same collection tube, to ensure an enrichment of plasmablasts/plasma cells (PB/PCs) in the sorted sample. Cells were collected in a BSA-coated Eppendorf tube filled with an RPMI + 10% FCS solution to avoid excessive cell loss. At baseline, first 10,000 total CD19+ B cells were sorted, and after adjusting the sorting gate to exclusively sort PB/PCs, another 1,201 CD19+CD38++CD24+CD27+ PB/PCs were sorted. For subsequent visits, all available PB/PCs were sorted (11,051 cells). In total, we sorted 22,252 cells. Directly after the sort, samples were spun down and resuspended into the recommended volume (RPMI + 10% FCS). Further sample processing took place at the leiden genome technology center (LGTC) in the LUMC according to the 10x Genomics protocols (**Figure S1**).

### 10X CITE-Seq transcriptomics and scBCR-rep sequencing

Nearly 22,000 B cells enriched in PB/PCs from different time-points were processed into single cells in a Chromium Controller (10X Genomics) (**Figure S1**). During this process, individual cells are embedded in Gel Beads-in-Emulsion (GEMs) where all generated cDNAs share a common 10X oligonucleotide barcode. Reverse transcription PCR and library preparation were carried out under the Chromium Single Cell 3’ v3 protocol (10X Genomics) per manufacturer’s recommendations. After amplification of the cDNA, a 5’gene expression library and paired heavy and light chain library were generated from cDNA of the same cell using Chromium Single Cell *VDJ* reagent kit (scBCR-rep library; v1.1chemistry, 10X Genomics). After library preparation, quality control was performed using a bioanalyzer (Agilent 2100 Bioanalyzer; Agilent Technologies). The libraries were sequenced in the NovaSeq6000 sequencer (Illumina) using the v1.0 sequencing reagent kit (read length: 2 × 150bp).

### Unsupervised clustering of single cell transcriptomics data

The raw sequencing data from different visits were processed using Cell Ranger (v3.1.0) against the GRCh38 human reference genome with default parameters. Single-cell RNA-seq data were processed in R with Seurat (v3.2.3) [9]. Raw UMI count matrices for both expression and hashtags oligos (HTO) generated from the Cell Ranger 10X Genomics pipeline (https://www.10xgenomics.com/) were loaded into a single Seurat object. Cells were discarded if they met any of the following criteria: percentage of mitochondrial counts > 25; the number of unique features < 100 or >5000. Furthermore, mitochondrial genes, non-protein-coding genes, and genes expressed in fewer than 3 cells were discarded. Moreover, the immunoglobulin *V* genes were removed from the counts for clustering analysis. The scaled counts of the constant genes were added to the metadata of the Seurat object. We used MULTIseqDemux [10] with default parameters to demultiplex the data to independent visits (corresponding to HTO1-HTO5 tags) and eliminated doublets and negatives among the cells (**Figure S2**). We found a few T cells (∼20-30 cells) in our data which clustered separately and we removed them from further analysis.

After retaining 5010 singlets, we log normalized the gene counts for each cell using the ‘NormalizeData()’ function of Seurat with the default parameters. The normalized data were scaled again using ‘ScaleData()’. The G2M and S phase scores were regressed out from the Seurat object. Principal component analysis (PCA), selecting the 10 most varying principal components, and t-distributed stochastic neighbor embedding (tSNE) dimension reduction were performed. A nearest-neighbor graph using the 10 dimensions of the PCA reduction was calculated using ‘FindNeighbors()’, followed by clustering using ‘FindClusters()’ with a resolution of 0.6. FACS-like plots were generated using transcript average cell scoring implemented in ‘TACSplot()’ [11].

### Differential gene expression analysis

For comparisons between expression values, the Seurat function ‘FindMarkers()’ was used with the ‘Wilcox method’. Cell type markers were obtained using the ‘FindAllMarkers()’ function with a Wilcoxon signed-rank test. The fold changes of the mean expression level of genes between the selected cell populations were calculated. The pathway analysis of the relevant genes was performed using ‘ClueGO’ [12].

### Identification of *V(D)JC* genes in the reconstructed BCRs

Demultiplexing, gene quantification and BCR contig and clonotype assignment were performed using the Cell Ranger (v.3.0.2) *V(D)J* pipeline with GRCh38 as reference. In order to get the BCR of a single cell, we obtained high-confidence contig sequences for each cell and kept the contigs with productive BCR rearrangements. The assembled filtered BCRs from 6437 cells were submitted to IMGT HighV-Quest for the *V(D)J* annotation. The constant gene annotation of the BCRs was obtained from the Cell Ranger output. The usage of constant gene in this individual was identified from the single-cell data and was compared with the constant gene usage identified by the flow cytometry. All chains (single, pairs, threes or fours) were retained. Of 6437 cells with BCRs, 4051 single cells had one heavy chain recorded in the Seurat object. The repertoire analysis was performed on both the sets (6437 and 4051 cells) and the discussed set is implicitly mentioned in the text or figure legends. In case two or more heavy chains were recorded from one cell, both the chains were included in the repertoire analysis.

### Identification of B cell clonotypes

Multiple files obtained from the IMGT HighV-Quest were combined using an in-house Python script. The B cell clonotypes were identified with the same *V* family, *J* family, and 80% nucleotide identity in *IG* heavy chain CDR3 as computed by Levenshtein distance {Python}. As *V* gene assignment might be flawed by the assignment methods, we used the *V* family to call clonotypes which avoids inaccurate grouping of the clones in the first place. Also, a comparative assessment was performed of the clonotypes obtained when using *V* genes versus *V* family in assigning B cell clonotypes.

### Analysis of scBCR-rep and clonotypes

The Circos plots of *VJ* usage were made using circlize {R}. To plot identified clonotypes, *IGHC* usage, class-switching events and CDR3 length usage, ggplot2 {R} was used. We used bcRep {R} [13] to plot the amino acid usage in the amino acid sequence of the junction. BCellMA {R} [14] was used to plot the replacement amino acid mutations from IMGT HighV-Quest output tables. Clonal lineages were built using alakazam [15] and plotted using igraph {R}.

### Query tool to identify anti-toxoid BCRs

The currently profiled BCRs comprise of all the PB/PCs raised in response to the vaccine which comprises of multiple components. To identify the contribution of each component of the vaccine, we developed an in-house tool using Python. To show its functionality, junction sequences related to anti-TT, anti-DT, anti-FHA, anti-Prn and anti-PT were searched in the literature. We could only identify the CDR3 sequences for anti-TT, anti-DT and anti-PT. These sequences were searched in the Boostrix-specific BCR-rep obtained in this study. The search was not only performed based on the percentage identity, but a parameter of length flexibility was also incorporated; i.e. the search space will include junction sequences that are longer or shorter by a user defined length. The sequence similarity is calculated based on the Levenshtein distance of the junction sequences.

## Results

### Single-cell transcriptome landscape of the cells from a vaccinated individual

We measured the transcriptomics and BCR-rep data at the single-cell level from the longitudinal samples obtained from an individual following Boostrix vaccination. After applying quality checks to the transcriptomics data from 10,000 total CD19+ B cells and 12,252 CD19+CD38++CD24+CD27+ PB/PCs from five pre-defined time-points (**Figure S1**), we obtained 5,010 singlets (**Figure S2**). Unsupervised clustering using Seurat distinguished seven different cell clusters based on their transcription profile (**Figure 1A**). CD19, CD38, CD24 and CD27 markers used to define the cells by flow cytometry allowed discrimination between B cells (**Figure 1A; Clusters B1-B3**) and PB/PCs (**Figure 1A; Clusters P1-P4**). Based on the tag mapping we could identify the sampling time-point of individual cells. The highest number of cells was obtained from day 0 (d0) when both total B cell and PB/PCs were sorted. At follow-up visits, more PB/PCs were obtained at d5 and d7and clearly less at d10 and d14 (**Figures 1B, S2B**). This reflected the number of sorted cells (**Figure S1B**), but mostly post-vaccination fluctuations in the number of PB/PCs (**Figure S1C**), which were also observed in the flow cytometry data obtained from this donor previously [5].

**Figure 1:**
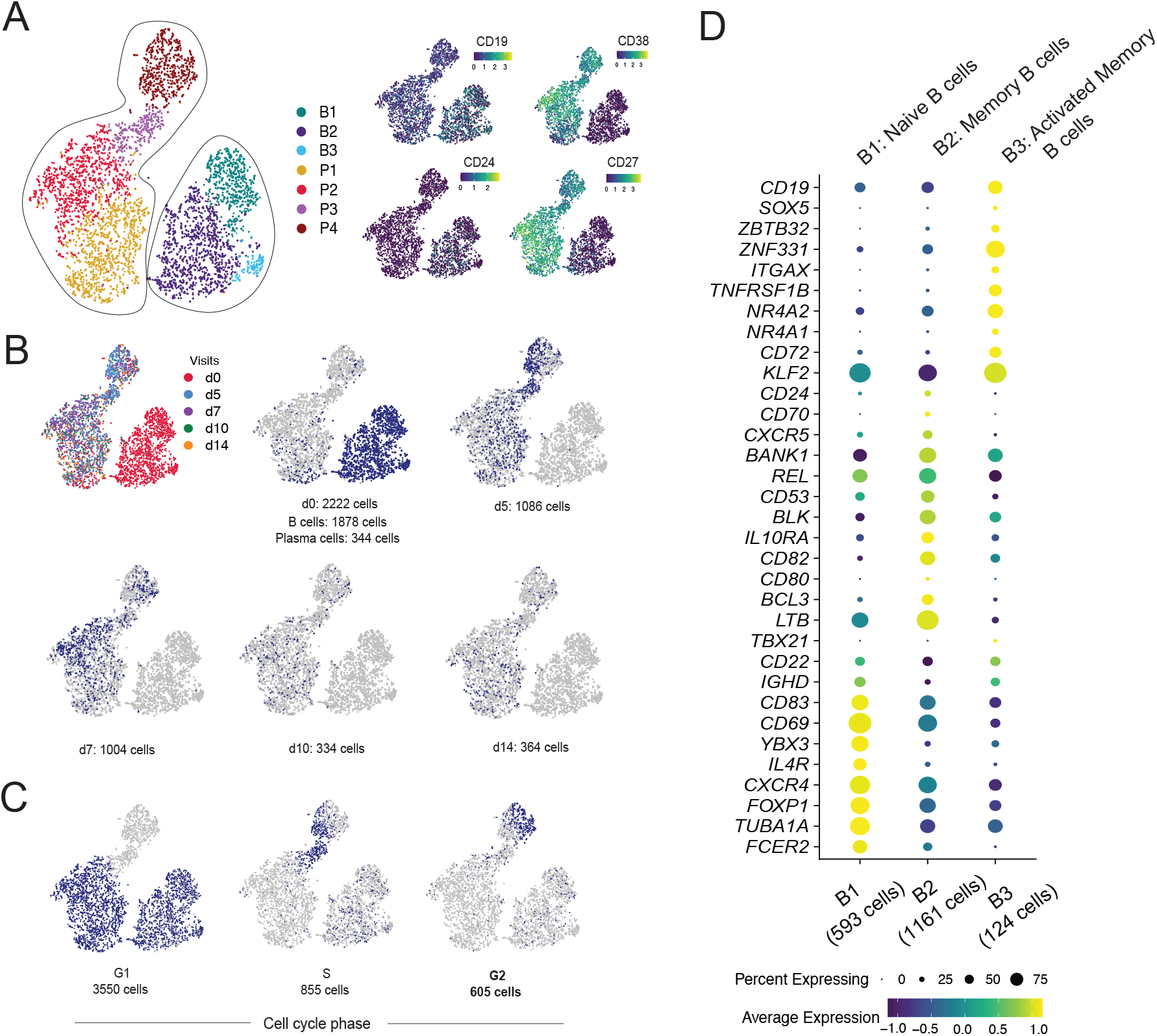
Identification of B-cell subpopulations by single cell-transcriptomics analysis. **A) The clusters of cells are represented on a tSNE plot based on the highly variable genes**. The clusters are named as B1-B3 for B cells and P1-P4 for PB/PCs clusters. The expression of the markers used for sorting the cells in the study is plotted on the tSNE plot. High expression of CD38 and CD27 marks the clusters P1-P4 as the PB/PCs. B**) Plotting tag information on the tSNE plot**. The cells are assigned to each visit and cells at d0 are assigned to B cells and PB/PCs based on their location in B1-B3 and P1-P4 clusters, respectively. **C) Visualizing cell cycle scores on tSNE plot**. G1, S and G2M cell-cycle scores are represented on the tSNE plot where P3 and P4 clusters comprise of S and G2M phase cells. **D) Dot plot of the gene markers used for identifying the B-cell subpopulations**. B1: naive B cells; B2: Memory B cells; B3: Activated memory B cells.

Despite having regressed out cell cycle influence, the P4 cluster separated based on the expression of the cell cycle genes (**Figure 1C)**. A majority of the cells in all the clusters, except P3 and P4, were in G1 with a few cells in S phase, whereas P3 and P4 clusters were heavily comprised of the cells belonging to S and G2M phase (**Figure 1C**), suggesting most proliferating cells are clustered together in P4. The cells present in G1 phase are the mature PB/PCs deemed to produce Abs which are mostly present in P1 and P2 clusters.

The B cell clusters i.e. B1-B3 were assigned individual identities based on the classical markers and/or literature. The B1 cluster expressed classical markers of naive B cells i.e. IGHD, *CXCR4*^*high*^ [16] and *FOXP1* [17] (**Figure 1D**). Cluster B2 was defined as memory B cells based on overexpressed of unique markers e.g. *BANK1, REL*. The genes *BCL3* and *LTB*, that promote Ab production, were also highly expressed in cluster B2 [18]. The *SOX5, ZBTB32* are markers for activated memory B cells which were also representative markers for cluster B3, hence the 124 cells in cluster B3 were identified as activated memory B cells [19]. The descending expression of *CXCR4* marker from naive to memory to activated memory B cells was observed in clusters B1 to B3, which was also mentioned in the similar subsets identified from an influenza vaccination study [19]. Overall, we could discriminate between naive B cells (cluster B1), memory B cells (cluster B2), activated memory B cells (cluster B3) and PB/PCs (clusters P1-P4) based on markers unique to these celltypes (**Figure 1A and 1D**).

### Flow cytometry guided plasma cell maturation in the single-cell transcriptome

Previously, we used longitudinal flow cytometry data from 10 vaccinated donors to establish how expression of selected cell surface markers changes during maturation of PB/PCs [5]. To visualize the plasma cell maturation in the here studied donor, the PB/PCs data from day 0, 5, 7, 10 and 14 were merged in the Infinicyt software (Infinicyt™ Software v2.0, Cytognos) and the maturation was defined by the downregulation of CD20 and upregulation of CD138 (**Figure 2A** left panel). The most mature cells exhibited CD20-CD138+ phenotype. Then we traced changes in expression of four additional markers i.e. CD19, CD27, CD62L, and CD38. Based on the expression of all six markers, we drew new maturation pathways and divided it into 6 continuous maturation stages (**Figure 2A** right panel).

**Figure 2:**
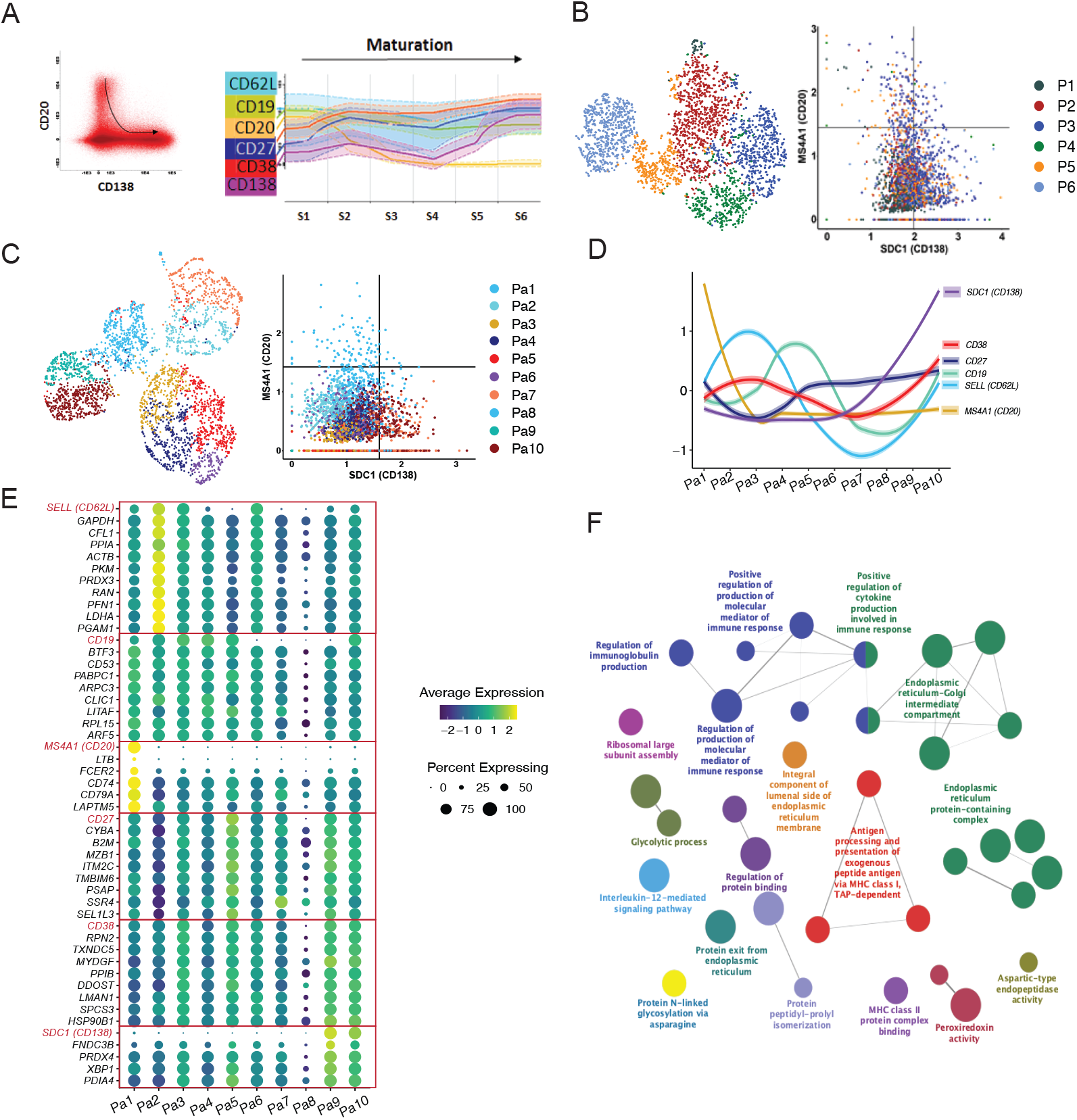
Plasmablast/plasma cell maturation using flow cytometry and single-cell transcriptomics data over time following vaccination. **A) Flow cytometric evaluation of cell surface markers in maturing** PB/PCs. Plasma cell maturation pathway was drawn in the Infinicyt software based on the surface expression of CD20 and CD138. The least mature PB/PCs were defined as CD20+CD138- and the most mature as CD20-CD138+. Left dot plot shows all PB/PCs from the same donor and time-points used in the sequencing, and black arrow indicates the direction of plasma cell maturation Each dot represents one cell. Subsequently, the entire maturation pathway was divided into 10 maturation stages (S1-S6), for which expression of additional cell surface markers was evaluated (right graph). **B) FACS-like plot to arrange plasma cell clusters P1-P6 in the maturation pathway defined by the expression of CD20 and CD138. C) Redefining stages of PB/PCs in single-cell transcriptome data using 6 markers pre-selected based on flow cytometry**. The PB/PCs were clustered using six pre-defined maturation markers and FACS-like plotting is used to stage the plasma cell clusters based on the maturation pathway. 10 clusters i.e. Pa1-Pa10 were obtained. **D) The spaghetti plot of all six markers visualized after staging the PB/PCs. E) Additional maturation markers identified using single-cell transcriptomics data based on the Pa1-Pa10 plasma cell clusters and corresponding stages**. Genes following similar expression patterns were identified using the TACSplot function. **F) The important pathways identified based on the top 20 similar genes to each of the six maturation genes**. The GO terms based on biological processes, molecular pathways, cellular components and immunological processes were merged. The pathways with <0.05 corrected P values were selected for visualization.

Similar to flow cytometry staging, after removing B cells, PB/PCs were clustered based on all the genes and arranged with phenotype ranging from CD20+CD138-(less mature) to CD20-CD138+ (most mature) (**Figure 2B**). However, the cells from different clusters were mixed and the maturation based on P1-P6 clusters could not be drawn. Therefore, we clustered the PB/PCs again based on the expression of only 6 genes (*SELL* (CD62L), *TNFRSF7* (CD27), *SDC1* (CD138), *ADPRC1* (CD38), *MS4A1* (CD20) and *B4* (CD19)) and used the expression of CD20 and CD138 to align the clusters with the maturation pathway of the PB/PCs. Herein, based on the maturation pathway defined by CD20 and CD138 markers, a trajectory from the blue colored clusters (Pa1) to the dark red ones (Pa10) can be drawn (**Figure 2C**). Based on the maturation of clustered cells the staging of the PB/PCs was performed from Pa1-Pa10 and the expression of other markers was assessed (**Figure 2D**).

Based on the 10 clusters/stages of the plasma cell maturation, we identified additional markers that might be changing during the maturation of the PB/PCs (**Figure 2E**). Genes with similar expression to the six maturation markers were selected from the single-cell transcriptomics data (**Table S1; Figure 2E**). The HLA locus specific genes were not included in the analysis. Several of these genes are involved in the regulation of immunoglobulin production, molecular mediation of immune responses, N-glycosylation, antigen processing and presentation and protein folding and cytokine production (**Figure 2F**). The majority of these pathways directly suggest the role of these genes in the Ig secretion, post-translational modifications to ensure correct folding of the proteins and henceforth an optimum response of the immune system to the vaccines. This is in line with the preparation for immunoglobulin secretion, which is the major role of most mature PB/PCs.

### Characteristics of *V(D)J* usage over time upon vaccination

Additional to transcriptomics profiling, we also measured the BCRs of the cells from the longitudinal samples obtained from an individual after Boostrix vaccination. The barcode mapped BCR information for each cell was obtained from the Cell Ranger output. We filtered out the unproductive rearrangements and could map at least one heavy or light chain for 4175 cells (**Figure S3A**). No chain was recorded for 835 cells (of which 818 cells were collected at d0) (**Figure S3A**), most likely owing to the low expression of BCR in non-plasma cells. We also observed that the cells comprised multiple chains varying from three to five heavy and/or light chains (∼8%). Therefore, we filtered out additional chains that were supported by a low number of reads as compared to the other chains from the same cell. In case multiple chains were supported by similar number of reads, we included them all in our analysis. After filtering, we found varying chain numbers (1, 3 or 4 chains) mostly for B cells identified at d0 (**Figure S3A**), which partly could be related to the low expression of BCR or possible doublets. The BCRs with 3 or 4 chains comprised of completely different *V, D, J* and *C* genes for both heavy and light chains, which might be related to the doublets that could not be removed in the filtering steps or the production of multiple chains in B cells [20]. For further comparisons in the manuscript, the BCRs from d0 B cells and PB/PCs were considered independently and post-vaccination BCRs were compared with d0 PB/PCs. Also, the expression of BCRs in B cells for both heavy and light chains was lower as compared to plasma cells.

The single cells with paired heavy and light chain information revealed preferential pairing between *IGHV* genes and *IGKV1* and *IGKV3* gene families (**Figure S3B**). *IGLV* genes mostly belonged to the *IGLV*1, *IGLV2* and *IGLV3* gene families. Also, over time, *IGKV1, IGKV3, IGLV1, IGLV2* and *IGLV3* gene families remained the most abundant, with minor variations in the usage which can be due to the limited number of cells or expansion of individual clones. However, no strong selection for a particular light chain was observed. On the contrary, we did observe changes in the usage of the *IGHV* genes over time (**Figure S3B**). Overall, the *IGHV3* was the most used gene family, however, the usage of *IGHV1, IGHV2, IGHV4* gene families increased in PB/PCs collected at d5 and d7 (around the peak of plasma cell response). To better understand the dynamics of the gene usage in *IGH* chains, we looked into the *V(D)J* gene usage. We found that *IGHD3* genes were used the most over all time-points, whereas the usage of *IGHV* and *IGHJ* genes changed over time (**Figure S3C**). We observed that the increased usage of *IGHV1, IGHV2, IGHV4* gene families was accompanied by an increased usage of *IGHJ2, IGHJ3* and *IGHJ6* genes post-vaccination.

As we observed increased usage of several *IGHV* and *IGHJ* gene families, we examined the usage of *IGHV* and *IGHJ* gene families in all cells across different visits at baseline and post-vaccination. We observed more restricted scBCR-rep in d0 PB/PCs as compared to d0 B cells (**Figure S3D**). While 55 unique of 158 *VJ* pairs were observed in d0 B cells, only 16 unique usage of 118 *VJ* pairs were present in d0 PB/PCs. The *IGHV3-23* and *IGHJ4* were the most used pairs in both d0 B cells and PB/PCs. The majority of the recombination events observed in d0 B cells used *IGHV1, IGHV3, IGHV4, IGHV5* families recombining with *IGHJ4*, however, it was highly selective in d0 PB/PCs, i.e. the *IGHV3* family genes and *IGHJ4* gene were used the most.

While comparing the PB/PCs from post-vaccination visits to d0 PB/PCs, we observed that the overall diversity increased at all the time-points post-vaccination (**Figure 3A**). We observed 50 and 65 new *VJ* pairs at d5 and d7 post-vaccination, respectively. Despite comparable numbers of sorted PBMCs and plasma cell counts at d10 and d14, we observed 24 new *VJ* pairs at d10 post-vaccination as compared to 35 new *VJ* pairs used at d14. Overall, we observed high usage of the *IGHV3-23* gene and the *IGHV4* gene family at all time-points including baseline. A clear expansion of *IGHV1* and *IGHV4* family genes especially *IGHV1-2, IGHV1-18, IGHV3-33, IGHV4-39, IGHV4-31* gene can be observed around the peak of plasma cell expansion (**Figure 3A**; d5, d7 and d10). *IGHV5-51* usage was also observed to be expanding at d10. The expansion of selected few *V(D)J* genes at the later time-points suggested the expansion of few clones with a delay. When comparing the relative usage of *IGHV* and *IGHJ* genes between visits post-vaccination, we observed that the usage of majority of the genes is comparable between visits with few exceptions, for example, the usage of *IGHV1-69* increased at d5 and d7 post-vaccination (**Figure 3B**). Furthermore, *IGHV1-2* and *IGHJ3* were used the most at d5 the most, whereas *IGHV1-2* and *IGHJ4* was used at d7, while at d10 and d14 the usage of *IGHV1-2* was minimal.

**Figure 3:**
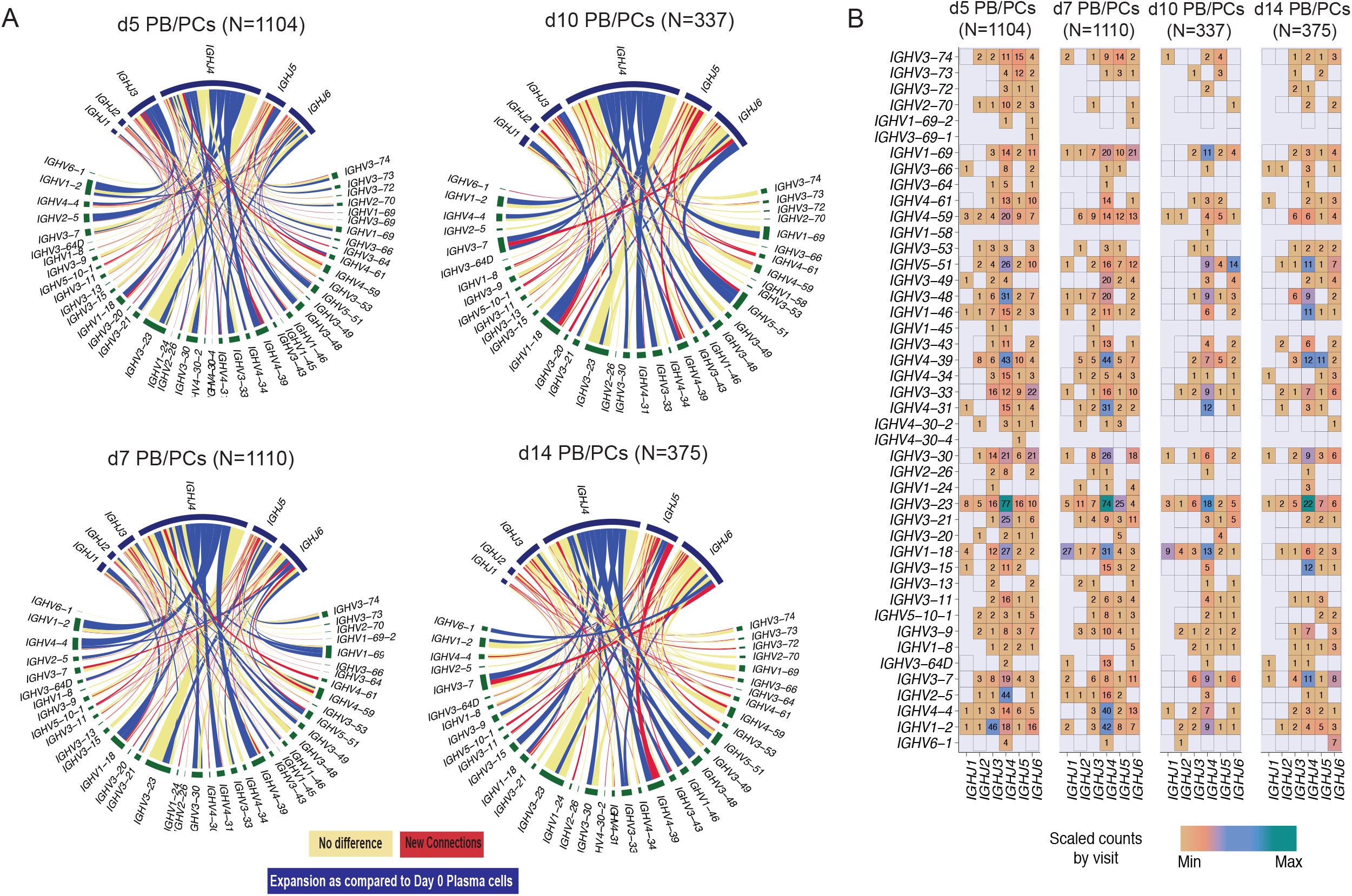
*VJ* gene usage in PB/PCs from all the post-vaccination visits relative to baseline. **A) CIRCOS plot of *VJ* gene usage in PB/PCs from all the post-vaccination visits relative to baseline**. Each ribbon represents a specific pairing between *V* and *J* genes, and the ribbon width represents the confidence bounds of the plotted data using loess method. Ribbons color shows no change (yellow) or expanded links (dark blue) as compared to d0 PB/PCs, or new recombination events (red). **B) Heatmap plot of *VJ* gene usage in PB/PCs from all the post-vaccination visits**.

### Constant gene usage over time upon vaccination

Assessment of the constant gene usage helps in understanding the raised effector function post-vaccination. To identify the usage of the constant genes in the profiled BCRs, we grouped the receptors based on the constant genes. *IGHM* was the most used constant chain in d0 B cells (**Figure 4A**), followed by *IGHD*. It was in line with the abundance of naive/pre-germinal center B cells in the pre-vaccination scBCR-rep. However, *IGHA1* was the most used constant gene in the d0 PB/PCs (**Figure 4A**). The composition of the usage of the constant chains changed after vaccination and *IGHG1* became the most used chain. The percentage of cells utilizing of *IGHA1* decreased at the peak of the plasma cell response i.e. d5 and d7, but at d10 and d14 returned to the levels comparable to baseline. The usage of *IGHD* was negligible in the PB/PCs post-vaccination. On the other hand, IGHM usage reduced at the peak of PB/PCs response which returned to original levels at d14. We observed a similar trend in the flowcytometry data, wherein the 67% of PB/PCs expanded at d7 was IgG1+ and the relative usage of IgA1 was reduced (**Figure 4B**). The IGHD expression was relatively low in the single-cell data limiting identification of the IgM+IgD+ B cells, which we could indeed identify in the flow cytometry data. With the paired chain information from single cells, we observed that the *IGKC* gene was the most used gene followed by *IGLC2* in all the post-vaccination visits. The high usage of the *IGKC* gene is in line to our previous results specifying high usage of *IGKV* gene families (**Figure 4C**). Interestingly, in d0 PB/PCs, *IGLC1, IGLC2* and *IGLC3* genes were used more frequently than other *IGLC* genes.

**Figure 4:**
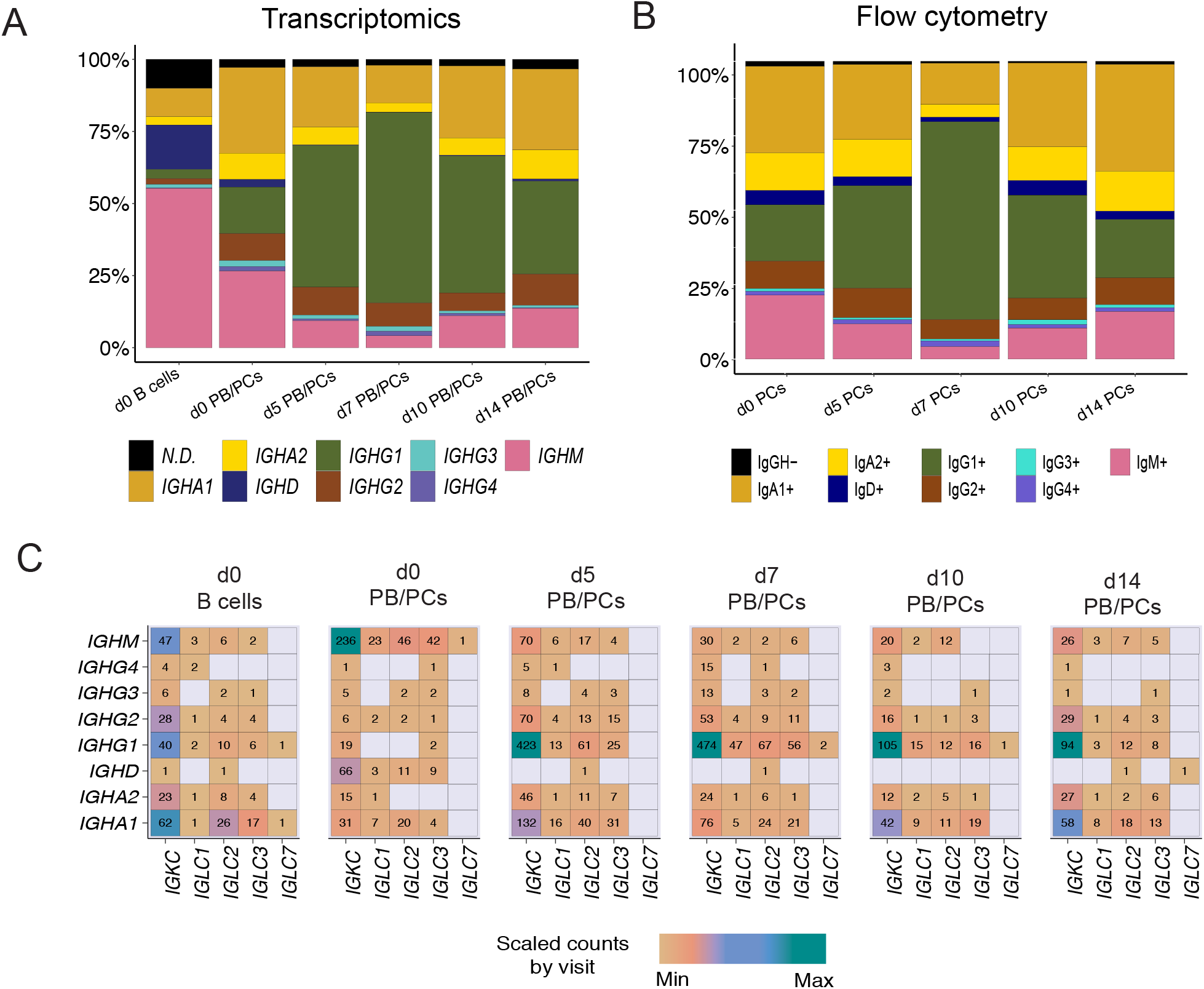
The dynamics of heavy and light constant gene usage in the BCRs over different time-points pre- and post-vaccination. **A) Usage of *IGH* constant genes in single-cell BCRs over time.** The PB/PCs and B cells at the d0 are considered separately. **B) Over time usage of constant genes in the PB/PCs as identified by flow cytometry (based on membrane and intracellular Ig expression). C) The usage of heavy and light constant genes across different visits**. The numbers in the heatmap are the number of chains using a given pair and the colors are ranged based on the scaled counts for each visit.

### BCR clonotypes over time upon vaccination

The BCRs were clustered into clones/clonotypes with a slight deviation from the classical definition of clonotypes; instead of genes, we used *V* and *J* families for grouping. Irrespective of the definition used, results of the clustering were highly similar (rand index of 0.99). The differences in the number of clusters were caused by the fact that several singletons clustered with other singletons with similar BCRs when the gene definition was relaxed by family (**Table S2**). Of 3334 clonotypes, 662 clonotypes could not be assigned any visit, 575 of which were singletons (8 clones with >10 BCRs; the largest clone comprises 129 BCRs). Overall, 936 of 3334 clonotypes had at least 2 cells. 369 of 936 clonotypes were shared between at least two visits of which 145 clonotypes had cells present on d0. For the 4051 cells derived from the known visits and with a rearranged productive heavy chain, **Figure 5A** shows the origin of the top 30 (449 BCRs) of 145 (2075 BCRs) clonotypes. The largest clone comprised of 46 B cells and was present in all visits but d14 (**Figure 5A**; C115). However, 30 additional B cells belonged to the same clonotype, but could not be assigned any visit (**Figure S4A**; the top second clone). The clonotype C115, consisting of only the *IGHG1* isotype, had an expansion of identical sequences across multiple visits including baseline as visualized by the largest pie in **Figure 5B**. This green node comprises of 58 individual BCR sequences wherein 34 of them are derived from d5 and 20 BCRs could not be assigned to any visit.

**Figure 5:**
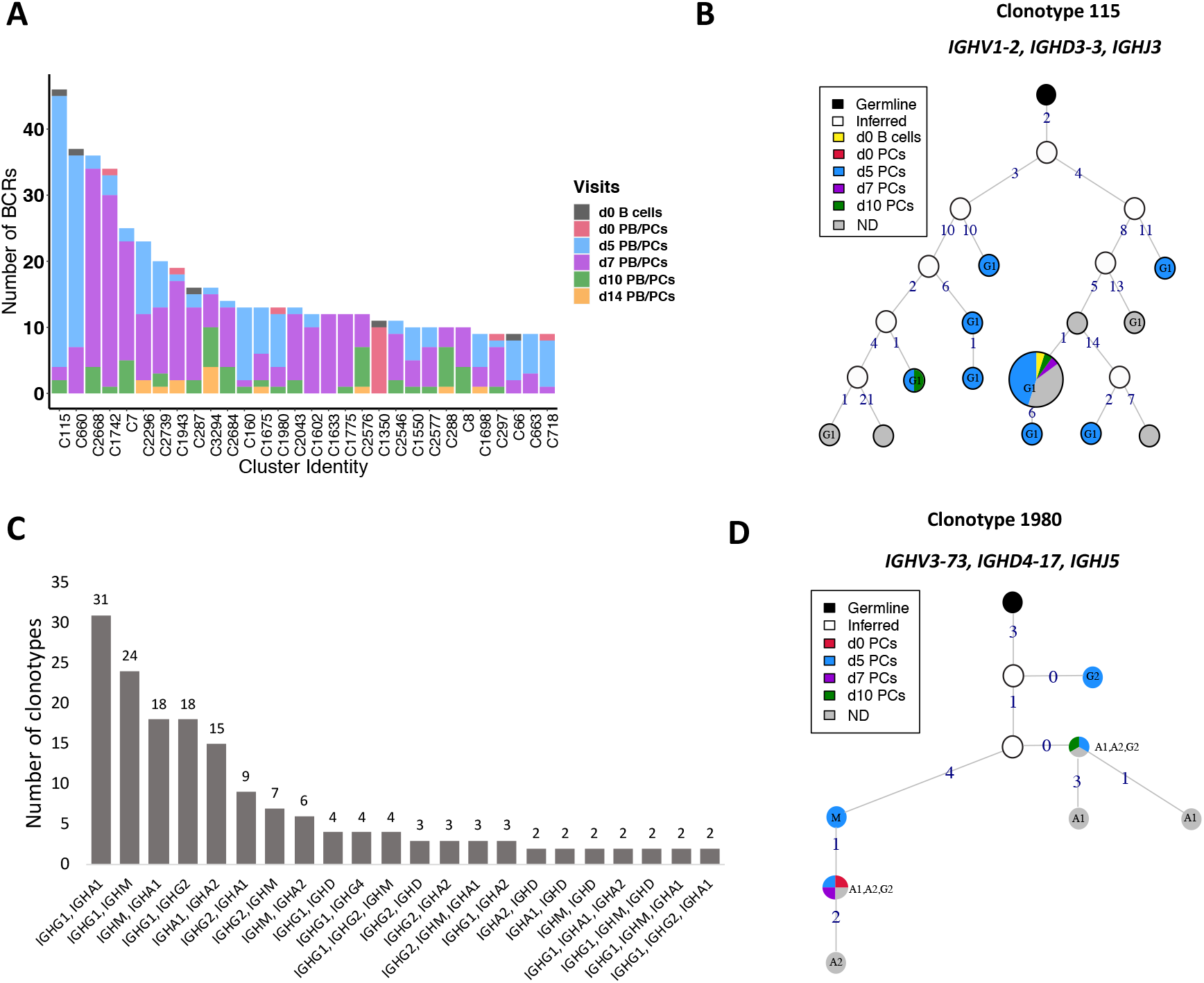
BCR clonotypes over different time-points. **A) 30 most numerous clonotypes over different time-points pre- and post-vaccination.** Contribution of different visits to the total size of a clone is indicated by colors. **B) Clonal lineage of clonotype 115**. The graph is constructed using *V* gene germline sequence as the root node. The size of each node is scaled and directly related to the number of BCRs for that particular sequence. The nodes are colored based on the visits from which sequences are derived. The unique sequences present across multiple visits are also grouped and colored differently and mentioned in the legend. Germline is colored black and the inferred nodes are colored white. The edge label indicates the number of different nucleotides as compared to the previous node. **C) A bar plot indicating the >2 clonotypes containing cells with multiple constant genes**. For example; each of 31 clonotypes consist of sequences using *IGHG1* and *IGHA1* constant genes. The number of clonotypes for each combination is mentioned on the top of bar. **D) Clonal lineage of clonotype 1980 with class-switch recombination events**. Additional inferred nodes were added to correctly represent the order of class-switching.

We investigated the clonotypes for the CSR events by querying clones with different *IGHC* chains. Of 369 clonotypes shared between different visits, 191 clonotypes had used the same constant gene and do not represent clonotypes with CSR. Interestingly, a majority of clonotypes (53%; 101 of 191) had only used the *IGHG1* gene, 21% clonotypes used *IGHA1* followed by 10% using *IGHM* and 8% using *IGHG2*. Thus, clonotypes followed the same trend as constant gene usage. Of 178 clonotypes that had multiple constant genes used by the BCRs, 17% had used *IGHG1* and *IGHA1* (**Figure 5C**). *IGHG1* is clustered with *IGHM* three times more (74%) as compared to *IGHG2* clustering with *IGHM*. Similarly, *IGHA1* is clustered with *IGHM* three times more than *IGHA2*, suggesting high CSR between *IGHM* and *IGHA1*. Overall, we found that 45% of these clonotypes contained *IGHM* and *IGHG* isotypes, followed by *IGHM* and *IGHA* (29%). The usage of multiple constant genes in the clonotypes suggest a high *IGHG1* response followed by *IGHA1*.

We selected a few of the large clonotypes to visualize the class-switching process via lineage construction. While some of the clones (e.g. clone 660, utilizing almost exclusively *IGHG1* with one *IGHA1*) were highly mutated and diverse (**Figure S4B**), others (e.g. clone 1980) showed a more conserved mutation pattern with very few mutations separating the clone members (**Figure 5D**). Clone 1980 is the 20^th^ largest clone where the *IGHA* and *IGHG* constant genes expanded with identical sequence in the *V* genes, which were only one to four mutations away from the *IGHM* harboring BCR (**Figure 5D**). This is an example of a receptor wherein the BCRs at later time-points harbor fewer mutations as compared to the germline than the BCRs sampled on d0. This suggests that this clone is derived from memory responses generated during previous antigen encounters.

Similarly, CDR3 length distribution was plotted for all the cells grouped by clonotype at baseline and each visit post-vaccination. We observed a larger proportion of clonotypes with CDR3 lengths of 12 amino acids at d5 (mean average length of CDR3: 15.8±3.8 amino acid residues) and 15 amino acids at d7 (mean average length of CDR3: 15.6±3.1 amino acid residues). We also observed a few clonotypes with longer junction region at d5 and d7 post-vaccination (**Figure S4C**).

### Mutation load in BCRs upon vaccination

The selection process for higher affinity Abs during the GC response also leaves an imprint on the *IGHV* genes, in terms of the type of amino acid substitutions observed in the cells which have successfully passed selection. Analysis of such mutation patterns at the nucleotide level allows us to understand the adaptability of the immune system to the antigens encountered during vaccination.

The majority of the BCRs in the activated memory B cells used the *IGHM* (34 cells) gene followed by the *IGHA1* (6 cells) constant genes. As expected, the mutation load between memory B cells (cluster B2) and activated memory B cells (cluster B3) were not significantly different. We observed that the *IGHG1* and *IGHM* BCRs in PB/PCs gained significantly more mutations (P < 0.05; <0.001) in the complete *V* gene as compared to the memory and activated memory B cells sampled at d0 (**Figure 6A**). The mutations in *V* region in genes using *IGHG1* constant gene are also accounted for in the CDR1 and FR3 region whereas for *IGHM* the significant differences were also observed in FR2, CDR2 and FR3 regions. We observed that *IGHV* of BCRs utilizing *IGHG1*, in general, harbored more mutations in post-vaccination visits as compared to other constant genes in all *V* regions (**Figure 6A**), which might be associated with the expansion in the *IGHG1* BCRs post-vaccination. Moreover, we found that the mutations in heavy chains were positively correlated with the mutations in the light chain in all the regions with significant P values (**Figure 6B**).

**Figure 6:**
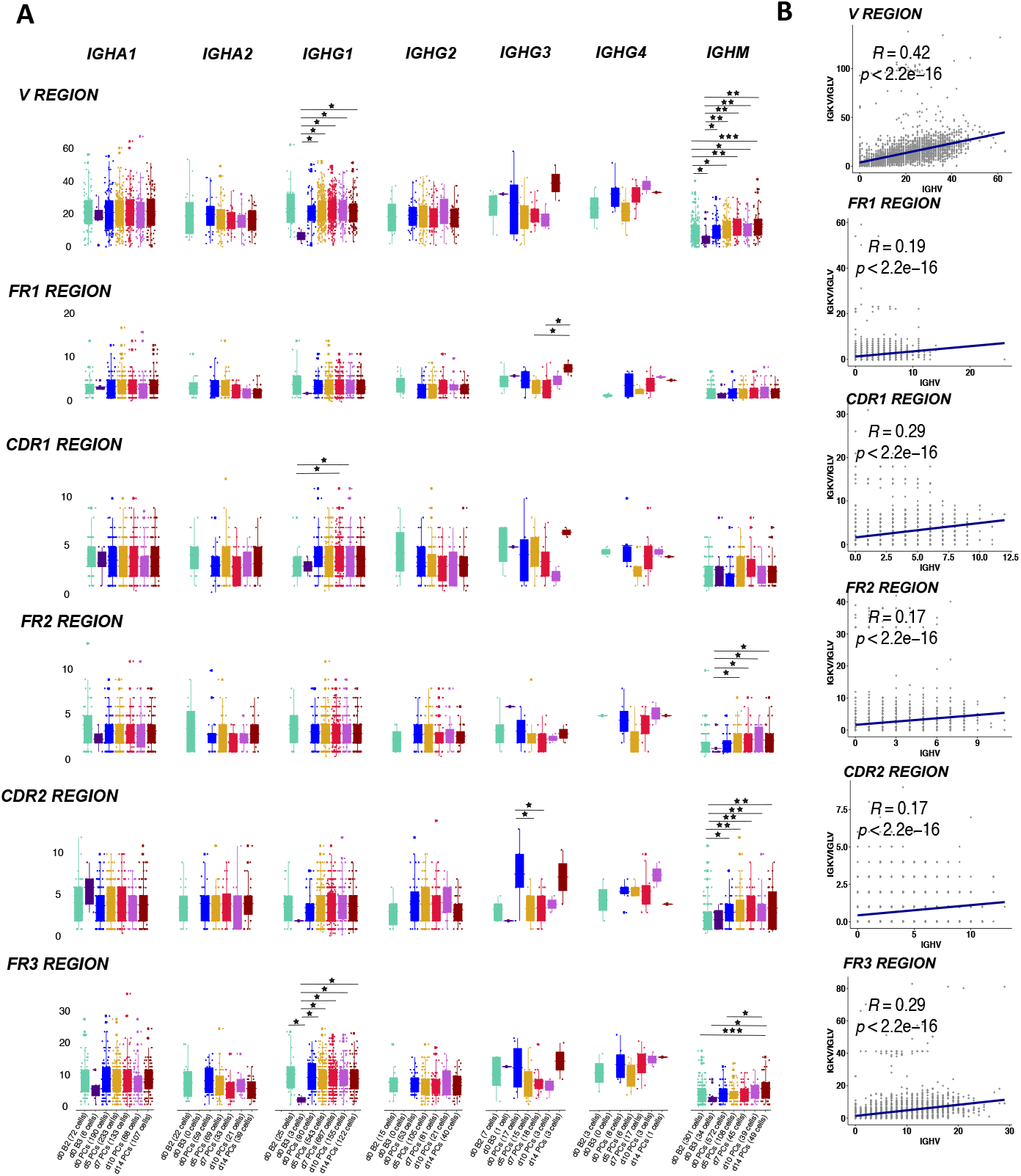
SHM level in the *V* genes in heavy and light chains. **A) The boxplot representing the number of mutations in the V genes of BCRs over different time-points pre- and post-vaccination in different constant genes.** The mutations have been identified in each cell over framework and complementarity determining regions in the *V* region. The cells at d0 are divided over naive B cells, memory B cells, activated memory B cells and PB/PCs. Each dot is a sequence from a single cell. The significant mean differences as calculated by the t test are highlighted by asterix as *: 0.05 - 0.01, **: 0.01 – 0.001 and *** < 0.001. **B) The correlation of mutation numbers in all the V regions between heavy and light variable genes for all cells with known paired constant BCRs**. The regression line is fitted and the correlation and p values are indicated on top.

To further confirm that the mutations in PB/PCs were antigen driven, we employed BASELINe [21] to detect the selection by analyzing mutation patterns in experimentally-derived Ig sequences. We observed a negative/neutral selection in framework regions and slightly positive selection in the CDR (**Figure S5**). The differences between the selection estimates in both FWR and CDR regions was highly significant (*P* = <0.03) except in FWR region at visit d10 (P = 0.25). The positive selection of CDRs and negative selection of FWRs correspond with normal antigen-specific B cells [22]. Moreover, we did not observe a varying trend i.e. increasing or decreasing selection pressure in CDRs over time suggesting that the selected receptors were already available at the beginning of the response and these were recruited over time for optimum immune response.

### BCRs specific to vaccine toxoids

To identify toxoid-specific lineages in the scBCR-rep, we searched literature for the CDR3 sequences known to be specific to antigens present in the Boostrix vaccine. We used the CDR3 amino acid sequences of the (mono-)clonal Abs for the anti-DT [23], anti-TT [24] and anti-PT [25] to query the related BCRs in our data (**Figure S6**). To account for individual-specific variability in the CDR3 sequences (as a result of SHM and affinity maturation processes specific to the individuals), we allowed a certain level of flexibility in our search. Apart from relaxing the identity of the CDR3 sequence at the amino acid level to 60-70%, the length of the CDR3 sequences was also allowed to vary by 1-3 amino acid residues.

We could not identify any relevant BCR sequence highly related to the anti-DT Ab sequences previously published. However, we identified 54 BCRs from 25 clonotypes that were highly similar to anti-TT Ab query sequences. 65% of the observed anti-TT BCR clones used the *IGHG1* constant gene followed by the *IGHA*1 (13%), *IGHM* (15%) and *IGHA2* (6%) genes. *IGHG1* was observed at all time-points, whereas *IGHA1, IGHA2* and *IGHM* were observed only until d7 post-vaccination. Clone 288 and 291 are 82% and 88% identical to the query anti-TT Ab sequence (**Figure 7A**). Similarly, clone 681 and 2083 are 80% and 86% identical to their respective anti-TT Ab query sequences. Clone 288 was the largest expanded clone with anti-TT specific BCRs (**Figure 7B**). This clone was present mostly at the later time-points post-vaccination. The presence of highly conserved BCR clones to the queried anti-TT Ab sequences suggests the BCRs can be highly conserved among different individuals. Furthermore, it also suggests that the immune response against TT is memorized efficiently by the immune system for secondary responses.

**Figure 7:**
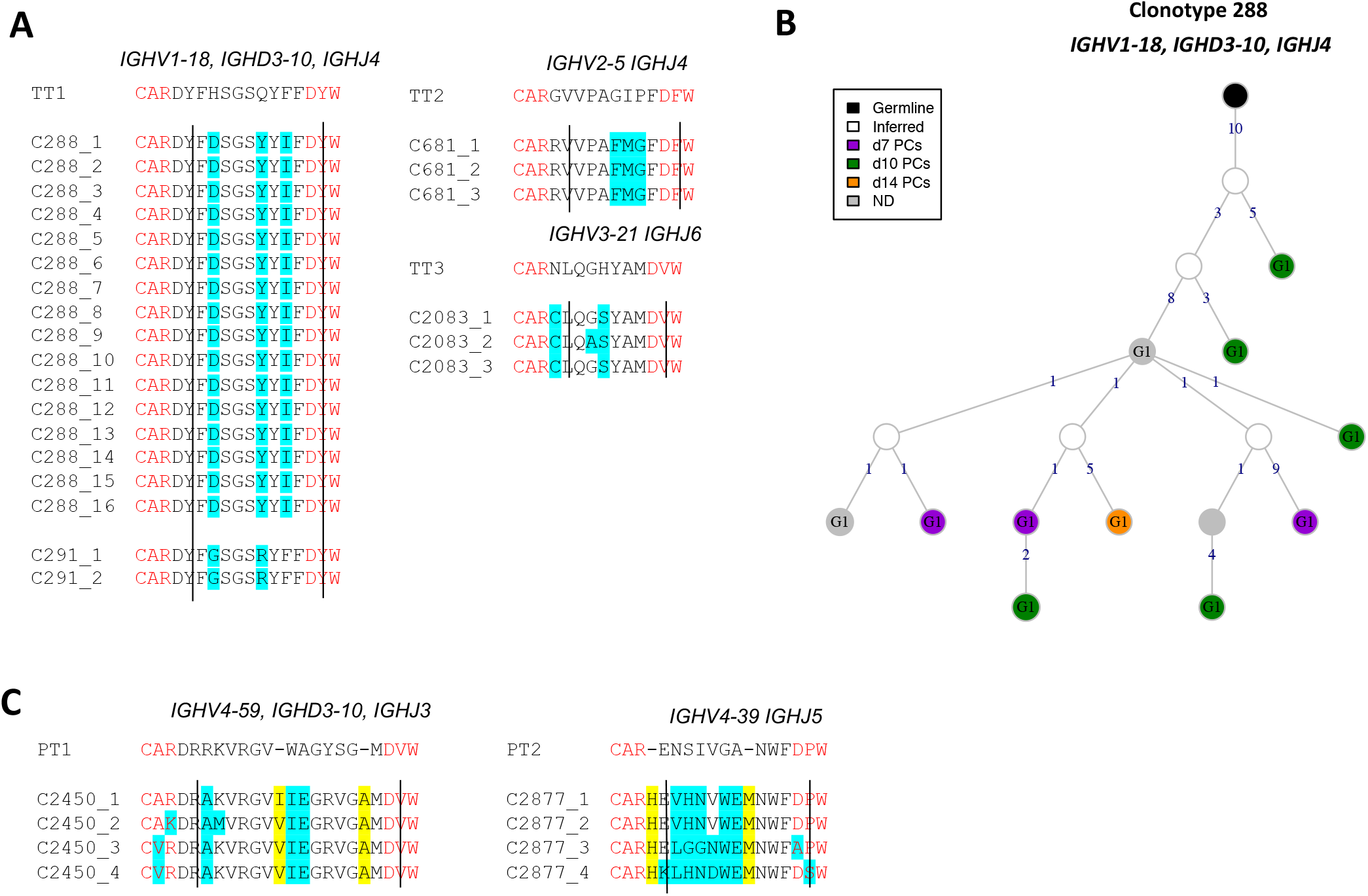
Toxoid-specific lineages upon Boostrix vaccination. **A) Highly conserved anti-tetanus toxoid specific BCR clonotypes.** Four clonotypes with >80% identity are aligned to the anti-TT Ab query sequences. The changed residues are highlighted in cyan and clonotype IDs are mentioned. The *V* and *J* gene residues are highlighted in red and demarcated by straight lines. **B) BCR lineage of anti-tetanus toxoid clonotype 288**. The largest anti-TT BCR clonotype is visualized. **C) Conserved anti-pertussis toxoid BCR clonotypes**. Two clonotypes with >70% identity are aligned to the respective anti-TT Ab query sequences. The length flexibility allowed the identification of these clonotypes. The changed residues are highlighted in cyan and the insertions are highlighted in yellow. Clonotype IDs are mentioned.

Analogously, we identified 20 BCRs that were similar to the anti-PT Ab query sequences. 55% of the hits used the *IGHG1* gene sampled from all visits, followed by 30% *IGHM* BCRs sampled from the memory B cells (cluster B3) at baseline. Unlike the anti-TT BCR sequences, we also observed a few BCRs using the *IGHD* constant gene. Two clones with maximum identity (∼70%) to the anti-PT Ab query sequence were found (**Figure 7C**). Both these clones used the *IGHG1* constant genes. Clone 2450 and 2877 are 73% and 71% identical to their respective anti-PT Ab query sequences. We did not observe highly conserved (>80%) CDR3 sequences to the anti-PT Ab query sequences suggesting either a unique individual-specific repertoire or recurrent adaptability of the immune response to the subsequent encounter of the antigen i.e. PT.

## Discussion

Understanding human B cell immune responses to infections is critical for vaccine development and assessment [26]. The repertoire of antigen receptor sequences changes during an encounter with foreign antigen from pathogens/vaccination [19,27]. Identifying the signatures of effective responses from the repertoire data is challenging and systematic analysis of the repertoire together with the dynamics of clonal lineages during infection/vaccination is crucial. Here, we systematically characterized the cell types in sorted B cells and PB/PCs using single-cell transcriptomics data and developed a robust workflow to capture the longitudinal changes in the BCRs and clonal lineages using repertoire data from the same cells. Moreover, we accorded our results with previously acquired flow cytometry data from the same individual [5].

The classical markers used in phenotyping by flow cytometry are not always captured in the sparse single-cell transcriptomics data which makes it challenging to define uniform populations in both the technologies. Therefore, the identification of naive B cells, memory B cells, activated memory B cells and plasma cell clusters was made using a combination of classical CD markers and other gene-based markers known in the literature. For example, *CXCR4*, a chemokine receptor, turned out to be an important marker for the identification of the naive B cells [16], whereas the classical markers *IGHD, CD19* were dimly expressed in this cell subset. Also, *CXCR4* is known to be progressively downregulated in transition from naive B cells to memory B cells to activated memory B cells [19], which we also observed in clusters B1, B2 and B3 (**Figure 1D**). The activated memory B cell state displays a hallmark of an effector B cell response and shares several markers with previously defined aged/autoimmunity-associated B cells, e.g. high expression of *ITGAX* (*CD11c*) [28]. Another activated memory B cell marker, i.e. *ZBTB32*, is known to modulate the duration of memory B cell recall responses in mice [29]. Defining the B cell populations accurately helps in better understanding of the molecular processes e.g. immunoglobulin production, antigen processing and presentation and glycolysation in vaccination/infection (**Figure 2F**), which in turn might be helpful in identifying therapeutic targets in pathogen immunity. Moreover, among all activated memory B cell markers, *ITGAX*, CD72 and *TNFRSF1B* are membrane markers, that in addition to the classical B cell markers can be used to sort activated memory B cells.

Despite regressing out cell cycle genes, we still observed a high score for the G2+S genes in the P4 cluster, suggestive of the most proliferating PB/PCs. Although we did not consider the cell cycle genes for lineage tracing, the P4 plasma cell cluster was the endpoint of the trajectory, which suggests the role of developmental genes other than cell cycle genes in the plasma cell development and proliferation. Interestingly, these P1-P4 clusters of PB/PCs do not accurately depict the developmental stages of PB/PCs as defined by the *CD20* and *CD138* markers in flow cytometry [5]. We showed that using flow cytometry data as a reference, the stages of plasma cell maturation could be tracked accurately and additional plasma cell maturation markers, contributing from the early to late plasma cell maturation pathway, were identified. For example, peroxiredoxin *PRDX3* (similar expression as *SELL* (*CD62L*)) suffices the metabolic requirements of PCs in early phase of development that are destined to secrete massive amounts of Ig [30]. *PPIA*, peptidyl-prolyl isomerase A, catalyzes the protein folding and might play an important role in folding the Abs in the early phase of plasma cell development. Additionally, *PABPC1* (expression similar to *CD19*) mediates the regulation of the switching of membrane IgH to secreted *IGH* isoform in PCs [31]. *CD74* (expression similar to *MS4A1* (*CD20*)) is directly involved in shaping the scBCR-rep by initiating a cascade that results in proliferation of B cells and rescues the cells from apoptosis [32,33]. *MZB1* (expression similar to *CD27*) is an important co-chaperone in plasma cell differentiation [34]. Loss of this gene can affect migratory properties of the Ab secreting cells and their trafficking and retention to the bone marrow [34]. Similarly, the *SSR4* gene plays an important role in protein assembly and trafficking [35]. Its high expression is related to IgG4-related diseases [36]. The markers *DDOST* and *PPIB* (expression similar to *CD38*) are involved in protein glycosylation and unfolded protein responses in the plasma cell development process as also identified in the pathway analysis. Finally, *XBP1* (similar expression as *SDC1* (*CD138*)) is responsible for late events i.e. increased protein synthesis in plasma cell differentiation [37,38]. Herein, 32% of 104 genes were found to be membrane bound and hence can be used in flow cytometry to dissect additional stages of plasma cell maturation and/or sort specific populations e.g. Ab-secreting cells. A few of these markers e.g. CD74 [39] and CD22 [40] have already been used in quantitative flow cytometry studies to distinguish normal B cells to that of the B-cell related malignancies.

The repertoire in response to infection/vaccination has generally preferences for specific *V, D* or *J* genes. For example, anti-hemagglutinin clones have been shown to use the *IGHV1-69* gene [35,41] and *IGHV4-34* gene [19] in influenza vaccination/infection studies. However, the usage of the *IGHV3-7* gene increased after Hepatitis B vaccination [42]. Also, the abundance of the *IGHV4-59, IGHV4-39, IGHV3-23, IGHV3-53, IGH3-66, IGHV2-5*, and *IGHV2-70* genes was found in SARS-CoV2 infection studies [43–46]. Similarly, we observed an increased usage of the *IGHV3-23* gene on all visits post-vaccination; the usage of the *IGHV1-2, IGHV1-18, IGHV3-33, IGHV4-39, IGHV4-31* genes increased at d5, d7 and d10 post-vaccination; and *IGHV5-51* was highly used at d10 post-vaccination. We found that none of the specific classes of the *IGHV* genes have driven the overall response. Moreover, we identified high usage of the *IGHD3* family and the *IGHJ6* gene, which is in general abundantly used in the general repertoire [47–49], most likely owing to the perfect recombination signal sequences [50,51]. In the light chains, we also observed a high usage of the *IGK* genes as compared to the *IGL* loci gene, reflecting the normal 3:2 ratio observed in humans at baseline [52–54], which was skewed to a ∼8:3 ratio at d5, suggesting selection mechanisms due to the antigen exposure.

We identified ∼900 of ∼3500 clonotypes with >1 BCR clustered together. The majority of them were present on d5 and d7 post-vaccination. Although, the PB/PCs expanded after vaccination are enriched for vaccine-specificity, we cannot rule out that some of the PCs may have a different specificity [55,56]. Approximately, 70% of the Ab-secreting cells in the influenza vaccination study have been shown to be influenza-vaccine specific, previously [57]. The clonotype classification allowed us to identify class-switching events, the majority of which contained *IGHM* and *IGHG*, followed by *IGHM* and *IGHA*. It is known from previous studies that ∼85% of the switches from IgM commonly comprises of IgG1, IgA1 and IgG2 in decreasing order [58]. Higher usage of the IgG subclasses (48% IgG1, 22% IgG2, and 9% IgG3) has been known previously from influenza vaccine responsive clones [19], which was also observed in our study. Altogether, the highest usage of the *IGHG1* gene and the highest number of clonotypes containing IgM and IgG suggest the vaccine target IgG-based immune response, common immune response in intramuscular vaccine administration.

The selection processes and molecular events affect the *V* gene of *IG* heavy chain differently than the light chains [59,60] resulting in less diversity in the light chain repertoire [61]. Although we observed significant correlations between the mutations in *V* regions in heavy and light chain genes, the correlation was < 45%. The high mutations in *IGHV* BCRs utilizing the *IGHM* class has been proposed as a mechanism of the immune system to maintain long-term memory to the vaccine [62]. Moreover, the high number of mutations in *IGHG1* and *IGHM* genes in CDR and FR3 regions might also be associated with the maturation of IgG affinity during the response to the Boostrix vaccine. As the vaccine is a booster and only one individual was used for the pilot study, it might be difficult to assess the role of booster in affinity maturation processes.

The dissection of the immune response to all the antigens and/or separate components of a vaccine, when cells are not pre-sorted based on their reactivity, can be helpful to estimate the response to each of the antigens. We identified a few clonotypes related to anti-TT and anti-PT BCRs from the repertoire, suggesting the public nature of the queried sequences. Identification of highly conserved anti-TT BCRs suggests an optimal repertoire for strong immunodominant epitopes. However, we observed that the anti-PT BCRs were not highly conserved which could be explained by either a unique private repertoire to the antigen or that each encounter with this antigen shapes the repertoire with additional mutations. The number of identified anti-TT and anti-PT clones are not directly related to the complexity of the proteins in the vaccine but related to the number of epitopes found in the literature. Additional support to this observation can be obtained with repertoire data obtained from antigen-specific B cells from multiple individuals.

Overall, we identified B cell populations and additional plasma cell maturation pathway biomarkers based on the transcriptomics data. Importantly, we used the longitudinal scBCR-rep data to understand the *V(D)JC* gene usage, clonal expansion and SHM events post-vaccination and their role in shaping immune responses. Together, we were able to dissect the immune responses to the components of the Boostrix vaccine.

## Acknowledgements

The authors gratefully acknowledge the Flow cytometry Core Facility at LUMC (coordinated by dr. K. Schepers, M. Hameetman, run by operators S. van de Pas, D. Lowie, J. Jansen, IJ. Reyneveld and former operators E. de Haas and G. de Roo) for their support. We are grateful to the subject, who volunteered to contribute to clinical study. We would also like to acknowledge Rick J. Groenland and Bas de Mooij, who helped in acquiring and sorting cells for the study. We would like to acknowledge Susan Kloet and the personnel from the Leiden genome technology center (LGTC) at the LUMC for providing the sequencing support. Finally, the authors would like to thank the team of Liesbeth Oosten and Jaap-Jan Zwaginga from the Dept. of Heamtology, LUMC, for their help with the clinical part of this study.

## Authors contributions statement

IK, AMD, EBvdA, MJTR, JJMvD and MAB: concept and design of the study.

AMD and MAB: Processed samples for single-cell sequencing and organized clinical part of the study.

IK: Analyzed the data.

IK and MAB: Interpreted the results.

All authors: Wrote the Manuscript, revisions and approval of the submitted version.

## Availability of data and codes

Raw and processed data have been deposited at Gene Expression Omnibus and are available via GEO accession GSE185427. The codes are made available via the GitHub repository https://github.com/InduKhatri/Single-cell-BCR.

## Supplementary data

**Figure S1: Overview of the time-points and the single-cell sorting strategy of samples obtained pre and post-vaccination from a single individual. A)** One healthy female volunteer (age 28; primed with DTP in infancy with no history of pertussis infection or vaccination in the past 10 years) was vaccinated using Boostrix vaccine. Peripheral blood samples were collected at d0 (baseline), d5, d7, d10 and d14 after vaccination. **B**) Workflow to sort 10.000 baseline B cells from baseline, and all PB/PCs from baseline, d5, d7, d10 and d14. The collection Eppendorf tube was precoated overnight with a 5% BSA solution (airdried afterwards) to reduce adherence of cells to the plastic. B cells were defined as Zombie-CD19+ cells. PB/PCs were defined as Zombie-CD19+CD38++CD24+CD27+. Created on Biorender.com. **C) Total B and plasma cell/plasma blast numbers of this donor as determined by flow cytometry**. The dashed lines indicate the baseline reference values for age-matched individuals (as described by Blanco et al, 2018 [63]). Ab; antibody, mAb; monoclonal Ab, L/D; life/dead marker, d; days after Boostrix vaccination.

**Figure S2: Identification of singlets in the CITE-Seq single-cell data based on the expression of HTOs**. The low-quality cells with unique features <100 or >5000 features, or/and >25% mitochondrial content were filtered out. After quality filtering steps, MULTIseqDemux was used to identify the singlets and match the singlets to respective tags hence visits. A tSNE of the singlets based on the tag expression is shown to depict the separation between tags.

**Figure S3: The number of recorded *IG* gene rearrangements and the *V(D)J* usage across different visits. A) The tSNE plot of the cells colored based on the number of recorded *IG* gene rearrangements**. The bar graph is used to indicate the numbers of heavy and light chains retained after filtering per cell. **B) The usage of heavy and light variable genes across each visit**. The numbers in the heatmap are the number of chains each combination of genes and the colors are ranged based on the scaled counts for each visit. **C) The alluvial plots to depict the usage of *V(D)J* genes in heavy chains**. The counts of each *V(D)J* usage are scaled (min:0, max:1) for comparison. The width of strata represents higher usage of the gene family. **D) Circos plot of the *V* and *J* genes in the heavy chains at d0 B cells and PB/PCs**.

**Figure S4: BCR clonotype expansions over different time-points. A) Top 30 most numerous clonotypes (909 BCRs) identified from 6347 cells over different time-points pre- and post-vaccination**. The count of clones per visit is colored. N.D. represent the receptors with no recorded visits. **B) BCR clonal lineage of clonotype 660**. The top third clonotype 660 with more than 50 BCRs is represented. Each BCR in the clonotype uses *IGHG1* and is different from its neighbors by one to 18 mutations, represented as edge labels. **C) CDR3 length distribution of the BCR amino acid sequences over different time-points**. N.D. represent the receptors with no recorded visits.

**Figure S5: Quantification of antigen-driven selection over-time post vaccination in PB/PCs**. The top panel depicts the density plot of the probability distribution of selection in CDR and FWR region in PB/PCs at baseline and all time-points post-vaccination. Negative sigma value indicates negative selection, neutral at sigma zero while positive value indicates negative selection. The plot in lower panel is simplified view of sigma values plotted for each time-points in both CDR and FWR regions. In the lower plot, we can clearly observe a neutral selection in FWR region in d10 PCs.

**Figure S6: The workflow of the query tool to identify toxoid-specific lineages in the scBCR-rep data**. The input sequences are searched against the database of consensus sequences of each clonotype. A LD score was obtained for each query sequence against similar consensus sequences (same *V* family, same *J* family; with user-defined CDR3 length adjusted and identity). The LD score is used to finally compile the related consensus sequence and the clonotypes for each query sequence.

**Table S1: List of genes with similar expression to the 6 maturation genes**. The gene names highlighted in green are the membrane markers. The dark grey shading indicates no pathway was associated with the gene, whereas light grey shading indicates the absent pathways.

**Table S2: Number of clusters identified by different grouping criteria with *V* genes and family**. The table is divided into three blocks based on varying BCR counts in clusters based on gene or family definition. The first block shows the number of clusters that actually differed between gene and family definitions; the second block represent unique clusters, whereas the last block, highlighted in yellow, represent the clusters that stayed constant even if the definition changed. The adjusted rand index for both the clustering methods was 0.99 suggesting very similar clustering output between the two definitions.

