## Supplementary Figures and Tables for "Longitudinal dynamics of human B-cell response at single-cell level in response to Tdap vaccination"

### Figure S1

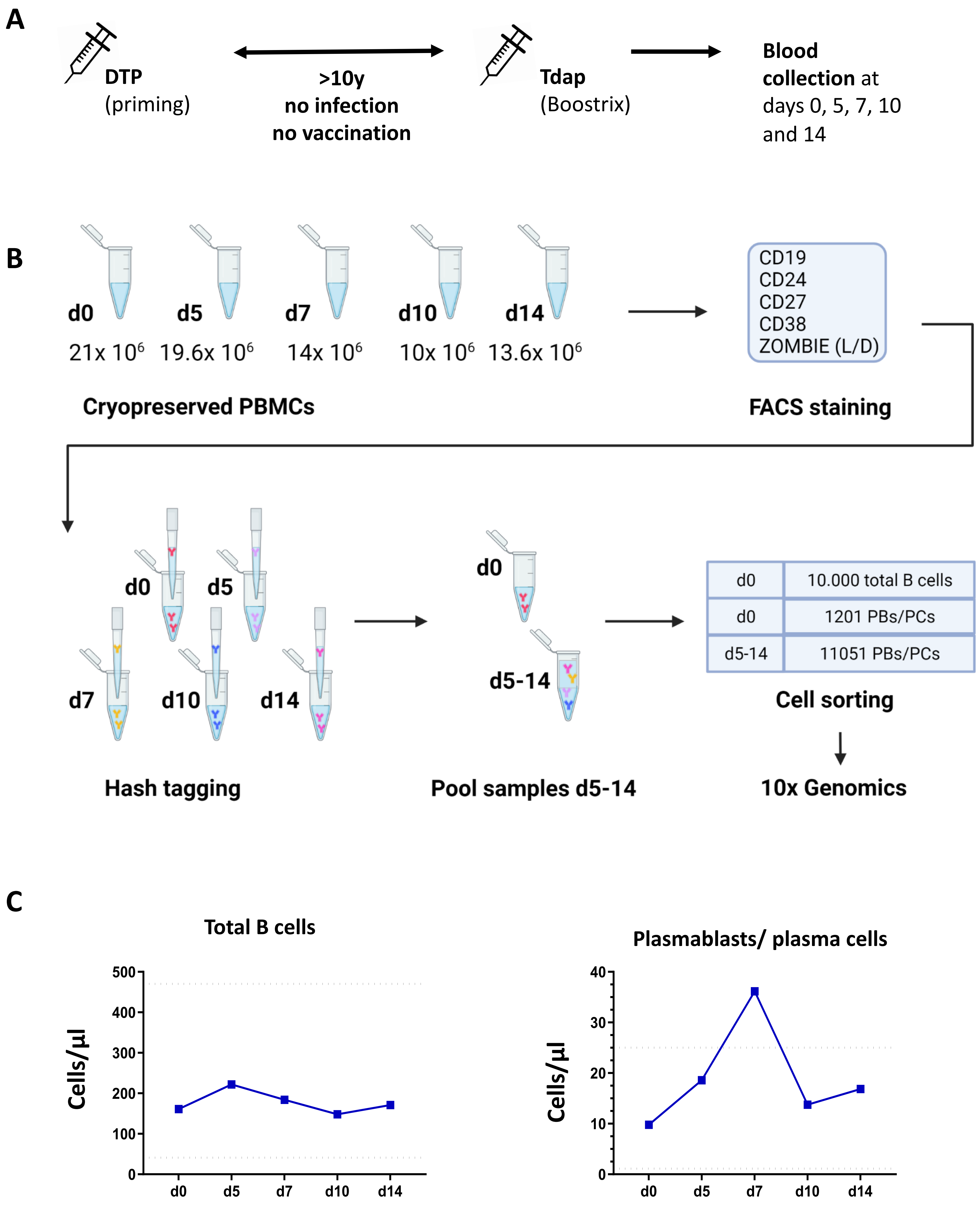
A
DTP
(priming)
>10y
no infection
no vaccination
Tdap
(Boostrix)
Blood
collection at
days 0, 5, 7, 10
and 14
B
d0
d5
d7
d10
d14
21x 10⁶
19.6x 10⁶
14x 10⁶
10x 10⁶
13.6x 10⁶
Cryopreserved PBMCs
CD19
CD24
CD27
CD38
ZOMBIE (L/D)
FACS staining
d0
d5
d7
d10
d14
Hash tagging
d0
d5-14
Pool samples d5-14
d0
10.000 total B cells
d0
1201 PBs/PCs
d5-14
11051 PBs/PCs
Cell sorting
10x Genomics
C
Total B cells
Cells/µl
500
400
300
200
100
0
d0
d5
d7
d10
d14
Plasmablasts/ plasma cells
Cells/µl
40
30
20
10
0
d0
d5
d7
d10
d14

Figure S2

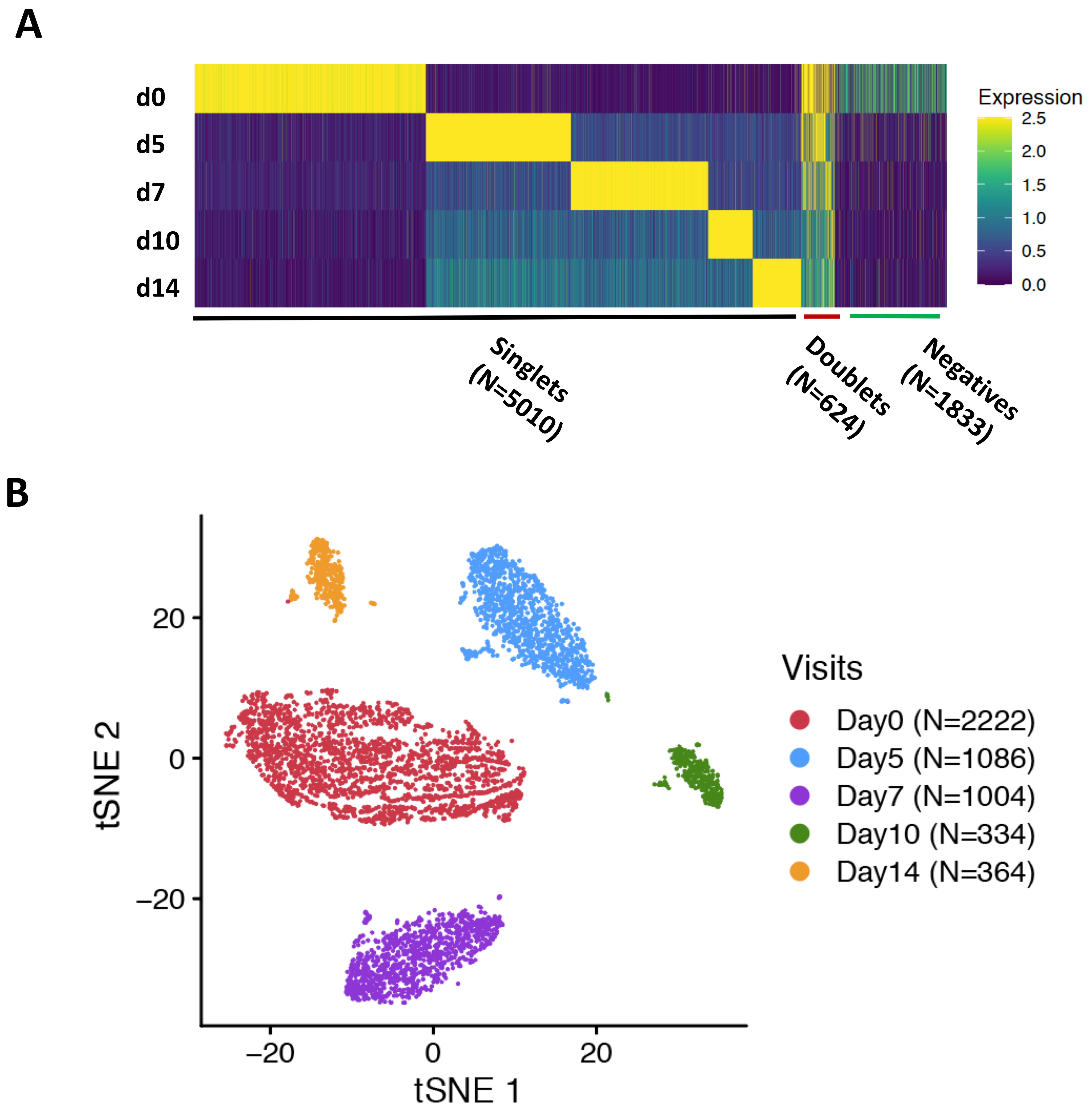
A
d0
d5
d7
d10
d14
Expression
2.5
2.0
1.5
1.0
0.5
0.0
Singlets (N=5010)
Doublets (N=624)
Negatives (N=1833)
B
tSNE 2
20
0
−20
−20
0
20
tSNE 1
Visits
Day0 (N=2222)
Day5 (N=1086)
Day7 (N=1004)
Day10 (N=334)
Day14 (N=364)

### Figure S3

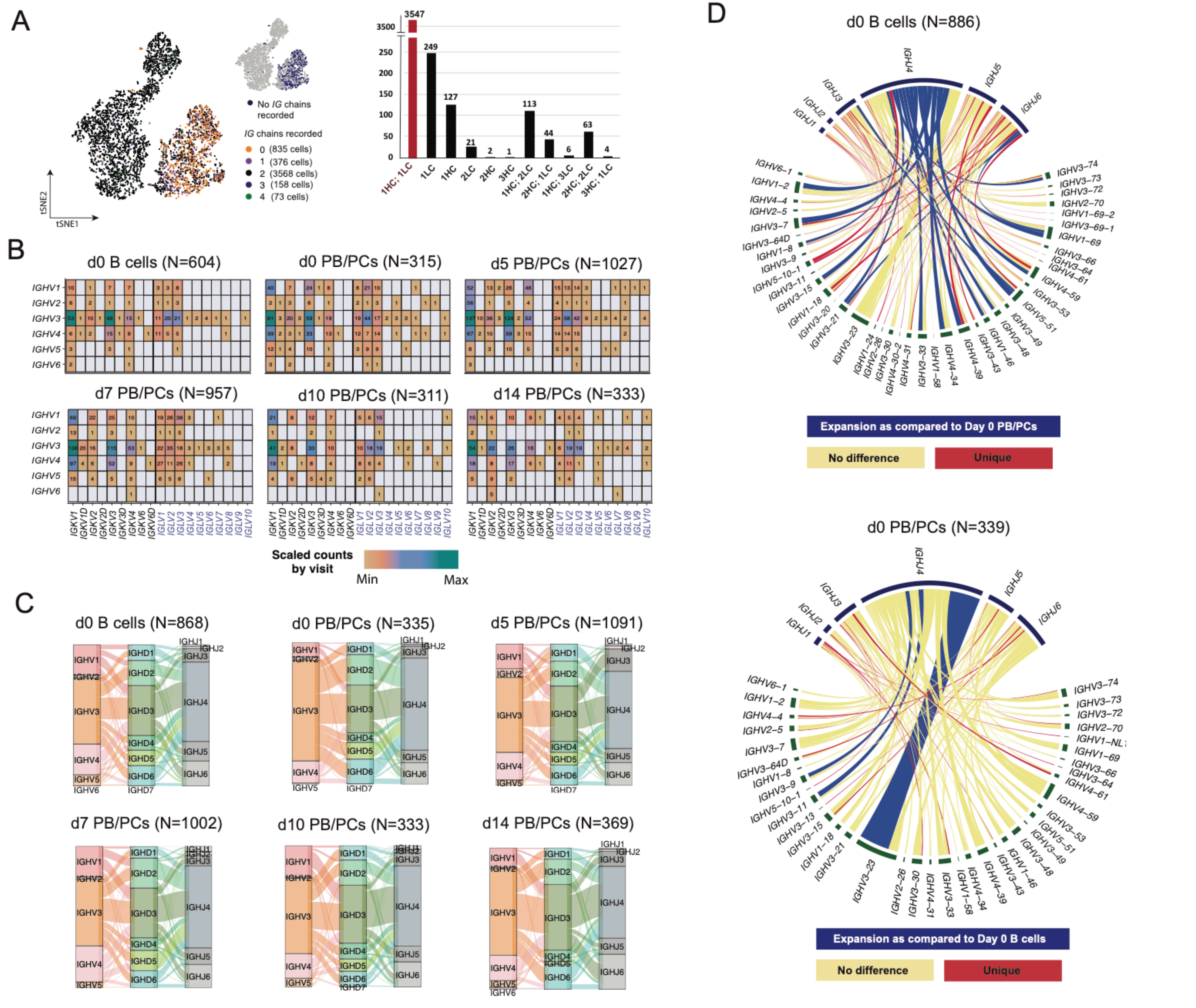

A
No IG chains recorded
IG chains recorded
0 (835 cells)
1 (376 cells)
2 (3568 cells)
3 (158 cells)
4 (73 cells)
tSNE1
tSNE2
3547
3500
249
127
21
2
1
113
44
6
63
4
250
200
150
100
50
0
1HC; 1LC
1LC
1HC
2LC
2HC
3HC
1HC; 2LC
2HC; 1LC
1HC; 3LC
2HC; 2LC
3HC; 1LC
B
d0 B cells (N=604)
d0 PB/PCs (N=315)
d5 PB/PCs (N=1027)
d7 PB/PCs (N=957)
d10 PB/PCs (N=311)
d14 PB/PCs (N=333)
IGHV1
IGHV2
IGHV3
IGHV4
IGHV5
IGHV6
IGKV1
IGKV1D
IGKV2
IGKV2D
IGKV3
IGKV3D
IGKV4
IGKV6
IGKV6D
IGLV1
IGLV2
IGLV3
IGLV4
IGLV5
IGLV6
IGLV7
IGLV8
IGLV9
IGLV10
Scaled counts by visit
Min
Max
C
d0 B cells (N=868)
d0 PB/PCs (N=335)
d5 PB/PCs (N=1091)
d7 PB/PCs (N=1002)
d10 PB/PCs (N=333)
d14 PB/PCs (N=369)
IGHV1
IGHV2
IGHV3
IGHV4
IGHV5
IGHV6
IGHD1
IGHD2
IGHD3
IGHD4
IGHD5
IGHD6
IGHD7
IGHJ1
IGHJ2
IGHJ3
IGHJ4
IGHJ5
IGHJ6
D
d0 B cells (N=886)
Expansion as compared to Day 0 PB/PCs
No difference
Unique
d0 PB/PCs (N=339)
Expansion as compared to Day 0 B cells
No difference
Unique

### Figure S4

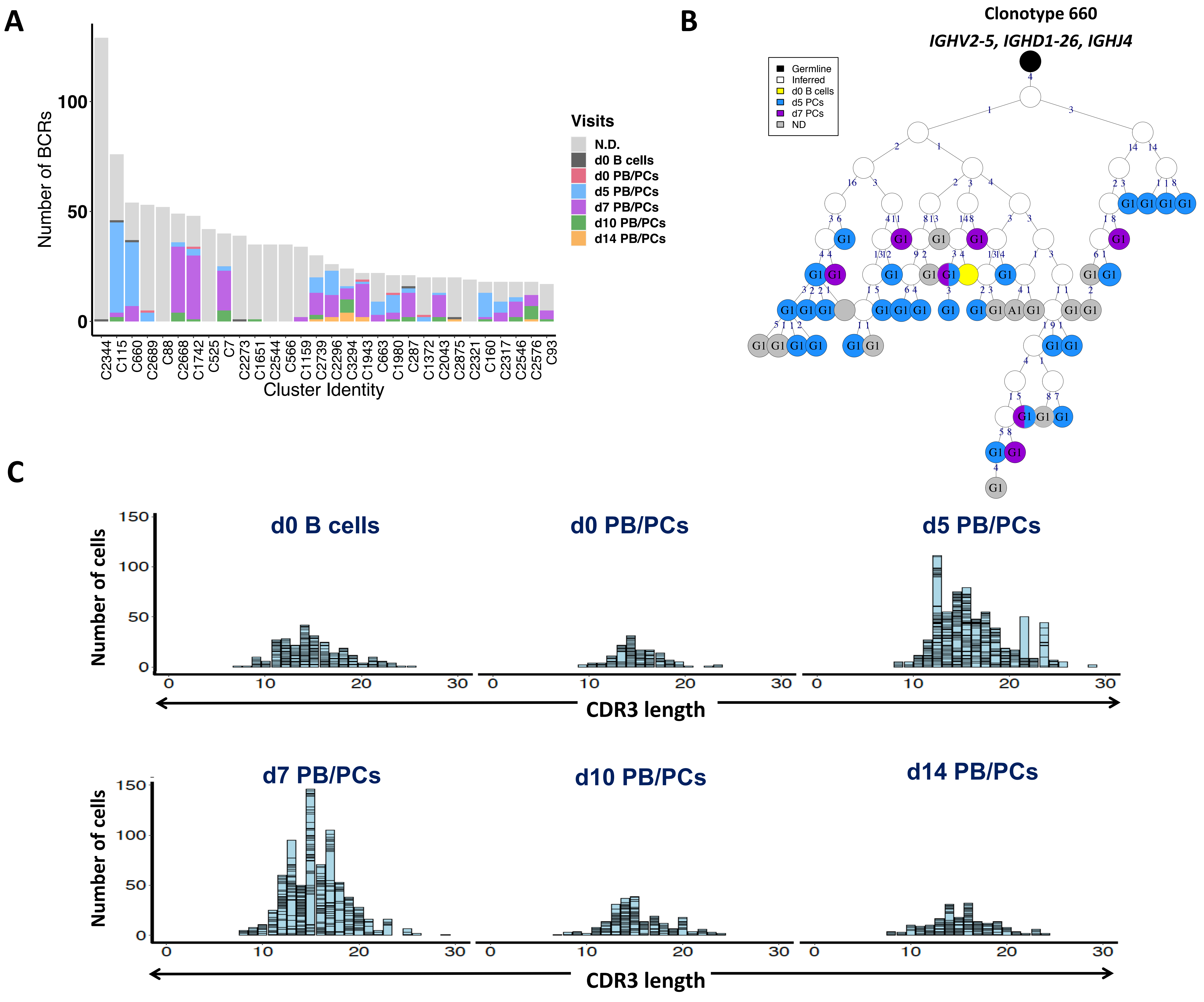
A
B
C
Number of BCRs
Cluster Identity
Visits
N.D.
d0 B cells
d0 PB/PCs
d5 PB/PCs
d7 PB/PCs
d10 PB/PCs
d14 PB/PCs
C2344
C115
C660
C2689
C88
C2668
C1742
C525
C7
C2273
C1651
C2544
C566
C1159
C2739
C2296
C3294
C1943
C663
C1980
C287
C1372
C2043
C2875
C2321
C160
C2317
C2546
C2576
C93
Clonotype 660
IGHV2-5, IGHD1-26, IGHJ4
Germline
Inferred
d0 B cells
d5 PCs
d7 PCs
ND
Number of cells
d0 B cells
d0 PB/PCs
d5 PB/PCs
d7 PB/PCs
d10 PB/PCs
d14 PB/PCs
CDR3 length
CDR3 length

**Figure S5**

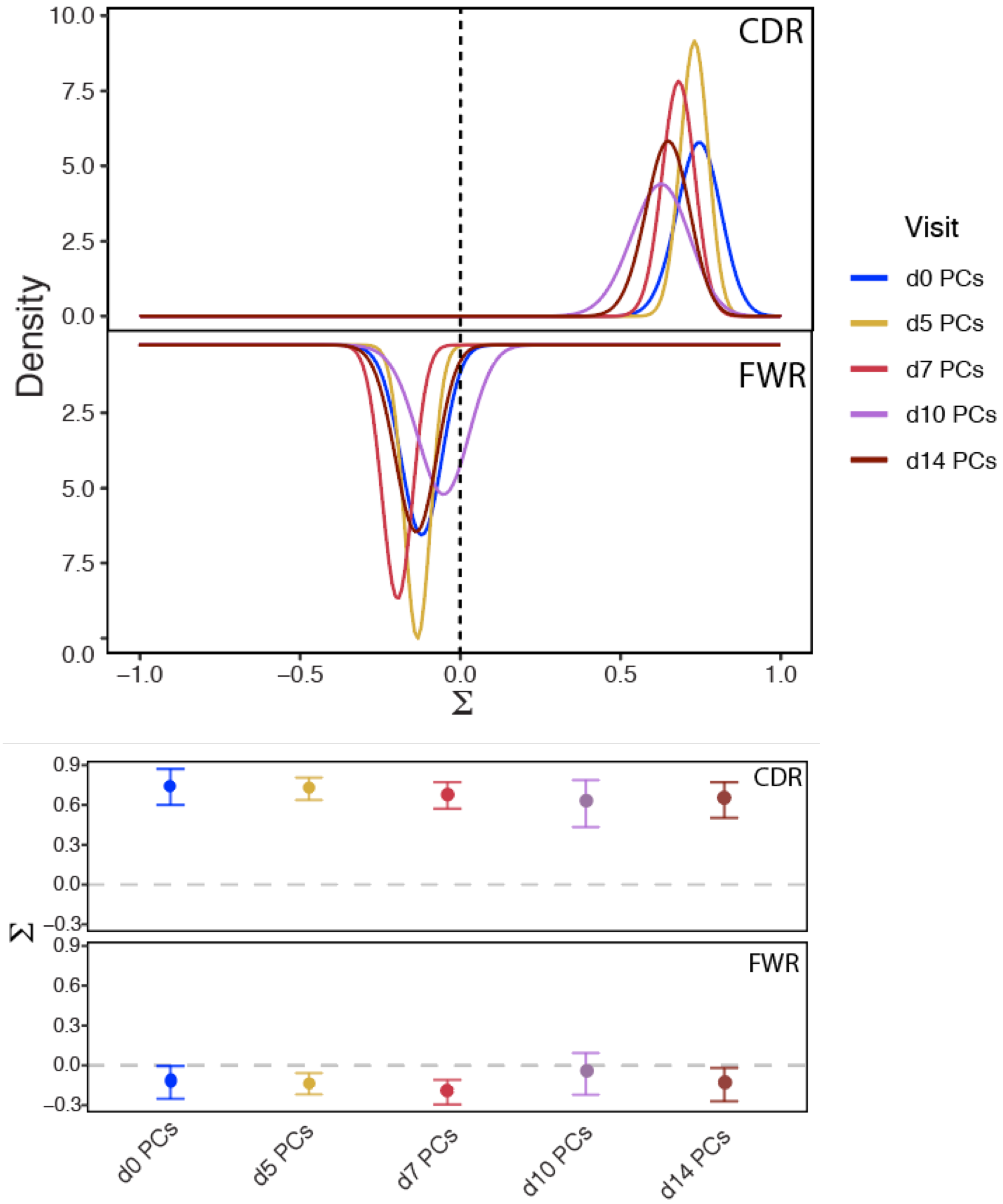
CDR
FWR
Density
10.0
7.5
5.0
2.5
0.0
−1.0
−0.5
0.0
0.5
1.0
Σ
Visit
d0 PCs
d5 PCs
d7 PCs
d10 PCs
d14 PCs
0.9
0.6
0.3
−0.3
d0 PCs
d5 PCs
d7 PCs
d10 PCs
d14 PCs

**Figure S6**

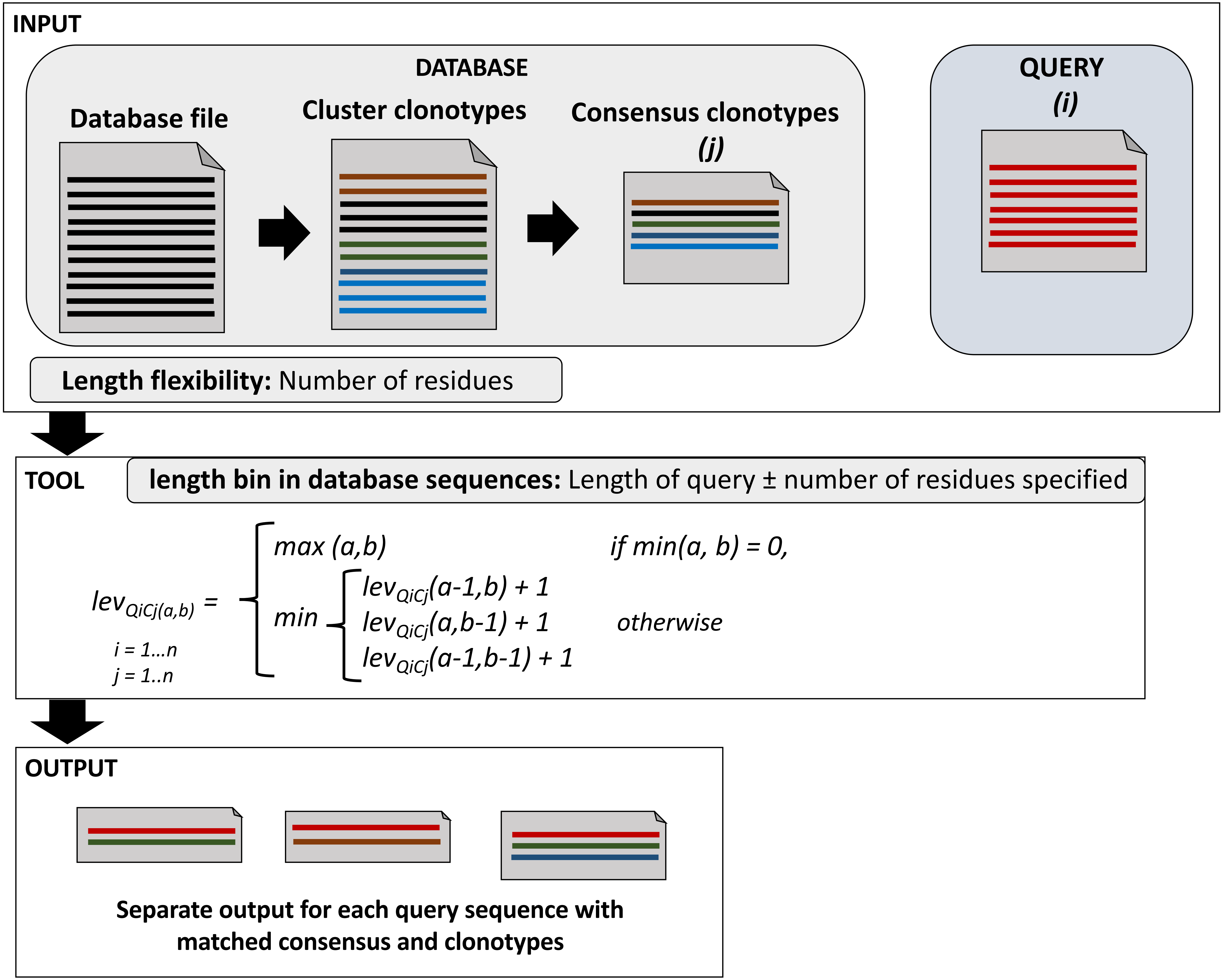
INPUT
DATABASE
Database file
Cluster clonotypes
Consensus clonotypes
(j)
QUERY
(i)
Length flexibility: Number of residues
TOOL
length bin in database sequences: Length of query ± number of residues specified
lev QiCj(a,b) =
max (a,b)
if min(a, b) = 0,
min
lev QiCj(a-1,b) + 1
lev QiCj(a,b-1) + 1
lev QiCj(a-1,b-1) + 1
otherwise
i = 1…n
j = 1..n
OUTPUT
Separate output for each query sequence with matched consensus and clonotypes

### Table S1

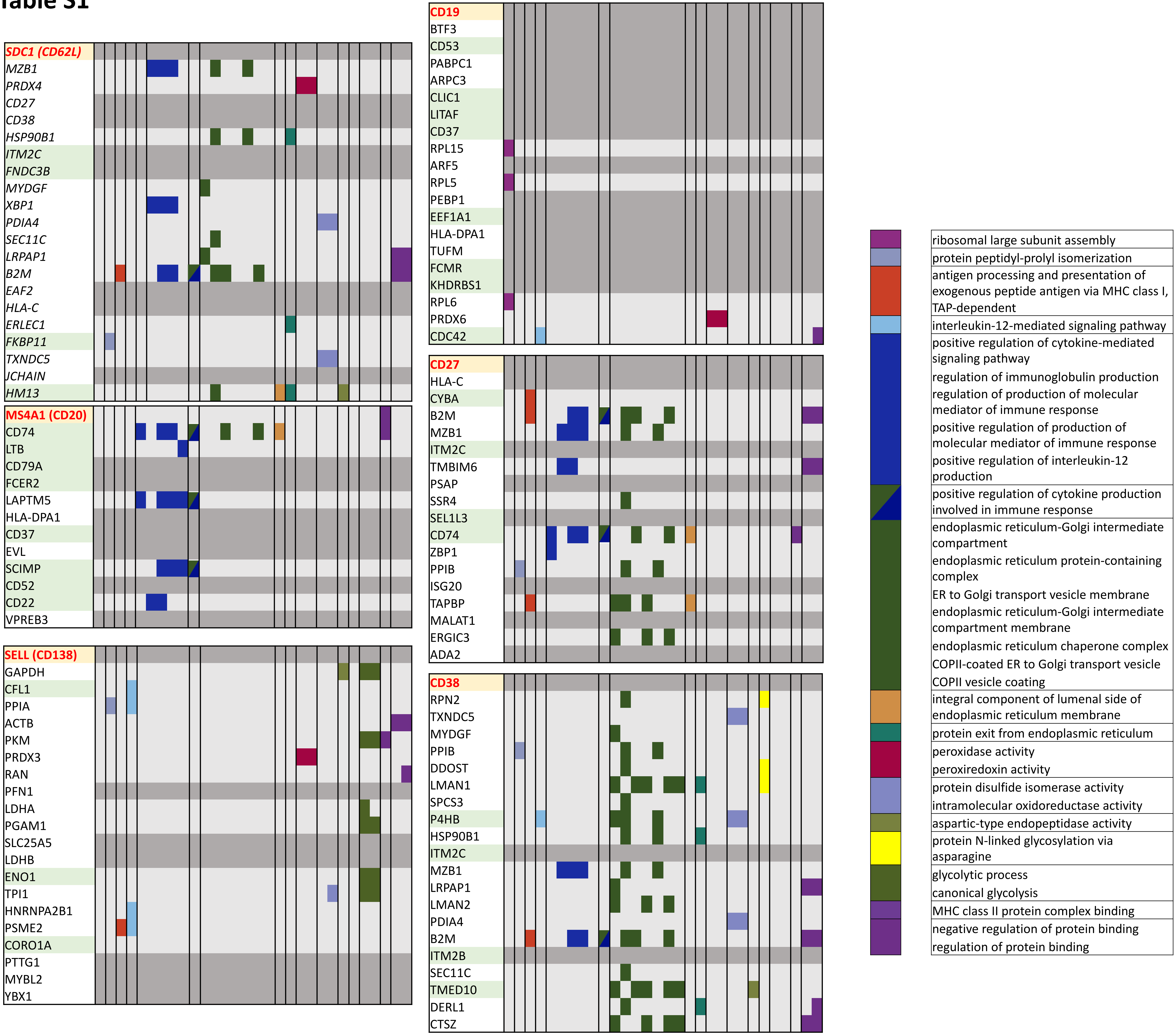

SDC1 (CD62L)
MZB1
PRDX4
CD27
CD38
HSP90B1
ITM2C
FNDC3B
MYDGF
XBP1
PDIA4
SEC11C
LRPAP1
B2M
EAF2
HLA-C
ERLEC1
FKBP11
TXNDC5
JCHAIN
HM13
MS4A1 (CD20)
CD74
LTB
CD79A
FCER2
LAPTM5
HLA-DPA1
CD37
EVL
SCIMP
CD52
CD22
VPREB3
SELL (CD138)
GAPDH
CFL1
PPIA
ACTB
PKM
PRDX3
RAN
PFN1
LDHA
PGAM1
SLC25A5
LDHB
ENO1
TPI1
HNRNPA2B1
PSME2
CORO1A
PTTG1
MYBL2
YBX1
CD19
BTF3
CD53
PABPC1
ARPC3
CLIC1
LITAF
CD37
RPL15
ARF5
RPL5
PEBP1
EEF1A1
HLA-DPA1
TUFM
FCMR
KHDRBS1
RPL6
PRDX6
CDC42
CD27
HLA-C
CYBA
B2M
MZB1
ITM2C
TMBIM6
PSAP
SSR4
SEL1L3
CD74
ZBP1
PPIB
ISG20
TAPBP
MALAT1
ERGIC3
ADA2
CD38
RPN2
TXNDC5
MYDGF
PPIB
DDOST
LMAN1
SPCS3
P4HB
HSP90B1
ITM2C
MZB1
LRPAP1
LMAN2
PDIA4
B2M
ITM2B
SEC11C
TMED10
DERL1
CTSZ
ribosomal large subunit assembly
protein peptidyl-prolyl isomerization
antigen processing and presentation of exogenous peptide antigen via MHC class I, TAP-dependent
interleukin-12-mediated signaling pathway
positive regulation of cytokine-mediated signaling pathway
regulation of immunoglobulin production
regulation of production of molecular mediator of immune response
positive regulation of production of molecular mediator of immune response
positive regulation of interleukin-12 production
positive regulation of cytokine production involved in immune response
endoplasmic reticulum-Golgi intermediate compartment
endoplasmic reticulum protein-containing complex
ER to Golgi transport vesicle membrane
endoplasmic reticulum-Golgi intermediate compartment membrane
endoplasmic reticulum chaperone complex
COPII-coated ER to Golgi transport vesicle
COPII vesicle coating
integral component of lumenal side of endoplasmic reticulum membrane
protein exit from endoplasmic reticulum
peroxidase activity
peroxiredoxin activity
protein disulfide isomerase activity
intramolecular oxidoreductase activity
aspartic-type endopeptidase activity
protein N-linked glycosylation via asparagine
glycolytic process
canonical glycolysis
MHC class II protein complex binding
negative regulation of protein binding
regulation of protein binding

### Table S2

| Number of unique BCRs | Number of clusters by $V_H$ Gene definition | Number of clusters by $V_H$ Family definition |
|---|---|---|
| 1 | 2747 | 2398 |
| 2 | 429 | 521 |
| 3 | 139 | 165 |
| 4 | 70 | 81 |
| 5 | 36 | 38 |
| 6 | 20 | 26 |
| 7 | 15 | 14 |
| 8 | 13 | 11 |
| 9 | 15 | 16 |
| 11 | 3 | 4 |
| 13 | 4 | 5 |
| 14 | 7 | 6 |
| 17 | 3 | 2 |
| 18 | 3 | 4 |
| 20 | 2 | 3 |
| 52 | 2 | 1 |
| 25 | 1 | |
| 28 | 1 | |
| 38 | 1 | |
| 75 | 1 | |
| 30 | | 1 |
| 39 | | 1 |
| 53 | | 1 |
| 76 | | 1 |
| 10 | 5 | 5 |
| 12 | 4 | 4 |
| 15 | 5 | 5 |
| 16 | 4 | 4 |
| 19 | 1 | 1 |
| 21 | 2 | 2 |
| 22 | 2 | 2 |
| 24 | 1 | 1 |
| 26 | 1 | 1 |
| 34 | 1 | 1 |
| 35 | 3 | 3 |
| 40 | 1 | 1 |
| 42 | 1 | 1 |
| 48 | 1 | 1 |
| 49 | 1 | 1 |
| 54 | 1 | 1 |
| 129 | 1 | 1 |
